# A Dynamic population of prophase CENP-C is required for meiotic chromosome segregation

**DOI:** 10.1101/2023.03.13.532437

**Authors:** Jessica E. Fellmeth, Janet Jang, Manisha Persaud, Hannah Sturm, Neha Changela, Aashka Parikh, Kim S. McKim

**Affiliations:** Waksman Institute and Department of Genetics, Rutgers, the State University of New Jersey, Piscataway, New Jersey, United States of America

## Abstract

The centromere is an epigenetic mark that is a loading site for the kinetochore during meiosis and mitosis. This mark is characterized by the H3 variant CENP-A, known as CID in *Drosophila*. In *Drosophila*, CENP-C is critical for maintaining CID at the centromeres and directly recruits outer kinetochore proteins after nuclear envelope break down. It is not known, however, if these two functions require the same CENP-C molecules. Furthermore, in *Drosophila* and many other metazoan oocytes, centromere maintenance and kinetochore assembly are separated by an extended prophase. Consistent with studies in mammals, CID is relatively stable and does not need to be expressed during prophase to remain at high levels in metaphase I of meiosis. Expression of CID during prophase can even be deleterious, causing ectopic localization to non-centromeric chromatin, abnormal meiosis and sterility. In contrast to CID, maintaining high levels of CENP-C requires expression during prophase. Confirming the importance of this loading, we found CENP-C prophase loading is required for multiple meiotic functions. In early meiotic prophase, CENP-C loading is required for sister centromere cohesion and centromere clustering. In late meiotic prophase, CENP-C loading is required to recruit kinetochore proteins. CENP-C is one of the few proteins identified in which expression during prophase is required for meiotic chromosome segregation. An implication of these results is that the failure to maintain recruitment of CENP-C during the extended prophase in oocytes would result in chromosome segregation errors in oocytes.

**Author Summary:** Meiosis in oocytes of diverse organisms, including humans and *Drosophila,* is characterized by a long prophase pause and a cell cycle arrest in meiosis I or meiosis II. These pauses could be a challenge for centromeres, whose components are replenished during G1, and then must remain with the chromosomes until the meiotic divisions. We have investigated the stability, prophase dynamics and function of centromere protein CENP-C. We show that CENP-C expression and loading onto centromeres during prophase is required for multiple meiotic functions. In contrast, the expression of other centromere partners CID/CENP-A and CAL1 is not required during meiotic prophase. Furthermore, expression of CID during prophase can be deleterious and result in ectopic kinetochore formation. CENP-C loading in prophase is required for sister centromere cohesion and kinetochore assembly. Our results provide the first description of CENP-C dynamics during meiosis and show that prophase expression is required for oocyte spindle assembly and function. CENP-C is among a small number of proteins that are required for the meiotic divisions but are loaded prior to prometaphase. Failure to maintain these proteins during the long prophase of oocyte meiosis may contribute to the increased aneuploidy associated with advanced maternal age.

## Introduction

The chromatin of the centromere is characterized by an H3 variant known as CENP-A rather than a specific sequence [1, 2]. One of its functions is to recruit kinetochore proteins for meiosis and mitosis [3]. Failure to maintain the centromere after DNA replication or failure to recruit an effective kinetochore during M-phase can result in chromosome segregation defects. In *Drosophila*, CENP-A is encoded by the *cid* gene [4]. CENP-A interacts with the “constitutive centromere associated network” (CCAN) complex in a wide variety of organisms such as humans and yeast [3]. The CCAN contains many proteins that form multiple links between the inner centromere and the outer kinetochore. In many vertebrate cells, two pathways defined by the components CENP-T and CENP-C provide the link between the centromere and outer kinetochore [5]. In some cases, one of these pathways has been lost during evolution. *C. elegans* and *Drosophila* are examples where most or all CCAN proteins have been lost and CENP-C forms the only linkage between the inner centromere and outer kinetochore.

Centromeres are maintained by loading new CENP-A once per cell cycle, but unlike replication-dependent histones, this does not occur during S-phase. Thus, CENP-A levels are reduced during S-phase and return to normal levels once during the cell cycle. The timing of CENP-A loading varies in different species and possibly cell types, but it usually occurs during M or G1 [6–8]. In yeast and mammals, maintenance of centromeric CENP-A depends on CENP-C, the Mis18 complex, and HJURP [9–15]. In *Drosophila*, the Mis18 complex and HJURP are absent, but CAL1 takes on a similar role by interacting with both CENP-C and CID [16–18]. CID, CENP-C, and CAL1 rely on each other for loading to maintain centromere identity [10, 16, 18, 19].

While CENP-C is required for centromere maintenance, it also has a direct role in kinetochore assembly by recruiting MIS12 in mammals [20–22] and *Drosophila* [23–25]. In many organisms, including *Drosophila* and mammals, the kinetochore is loaded once the nuclear envelope breaks down, and is only present during cell division. Although considered part of the inner kinetochore, CENP-C is present throughout the cell cycle. In *Drosophila,* no other kinetochore proteins except for MIS12 exhibit this behavior [26]. CENP-C recruits MIS12 [24, 25], and they directly interact to form an inner kinetochore complex that is present throughout the cell cycle [21, 23, 27].

Following the pachytene stage of meiosis, mammalian and *Drosophila* oocytes enter a long pause in prophase. If centromere proteins are loaded once per cell cycle, they would have to be maintained for a long time before kinetochores are assembled and cell division commences [28]. Cohesins, for example, are loaded during S-phase and deteriorate with age, causing aneuploidy in older mothers [29, 30]. CENP-A nucleosomes in mouse oocytes are stable and do not need to be maintained during meiotic prophase [28, 31]. Furthermore, CENP-C becomes immobilized at metaphase [32]. Thus, it is possible that the CENP-C loaded along with CENP-A in G1 is sufficient for kinetochore assembly. However, the stability of CENP-C during meiotic prophase has not been tested.

In this study, we investigated the meiotic prophase dynamics of CENP-C, CAL1, and CID in *Drosophila* oocytes. As with CENP-A in mouse oocytes [28], we found that CID is stable throughout *Drosophila* oocyte meiotic prophase, and there was little addition of new subunits. In fact, expression of CID during prophase can lead to ectopic kinetochore formation and abnormal chromosome segregation. CAL1 is gradually lost during prophase and is absent in metaphase I oocytes. CENP-C is unique because it is loaded at centromeres using an exchange mechanism during meiotic prophase, and this is required for assembly of the meiotic kinetochore. Thus, CENP-C is loaded independently of centromere maintenance in a process that is required for kinetochore assembly.

## Results

### CENP-C, but not CAL1 or CID, loads onto centromeres during prophase I

The analysis of centromere protein dynamics during prophase is facilitated by the organization of the *Drosophila* ovary. Each of the two *Drosophila* ovaries contains several strings, or ovarioles, of oocytes arranged in developmental order (Figure 1A). At the anterior end of each ovariole is the germarium, which includes mitotically dividing cells (region 1) and early meiotic prophase (regions 2-3). Region 3 (stage 1) oocytes leave the germarium and enter the vitellarium. Stages 1 to 13 last 5 days, during which meiosis remains in prophase while the oocyte grows and matures [33, 34]. At the posterior end of the ovariole (stage 13-14), the oocyte enters prometaphase. Meiosis arrests at metaphase I until passage through the oviduct and fertilization occurs.

**Figure 1:**
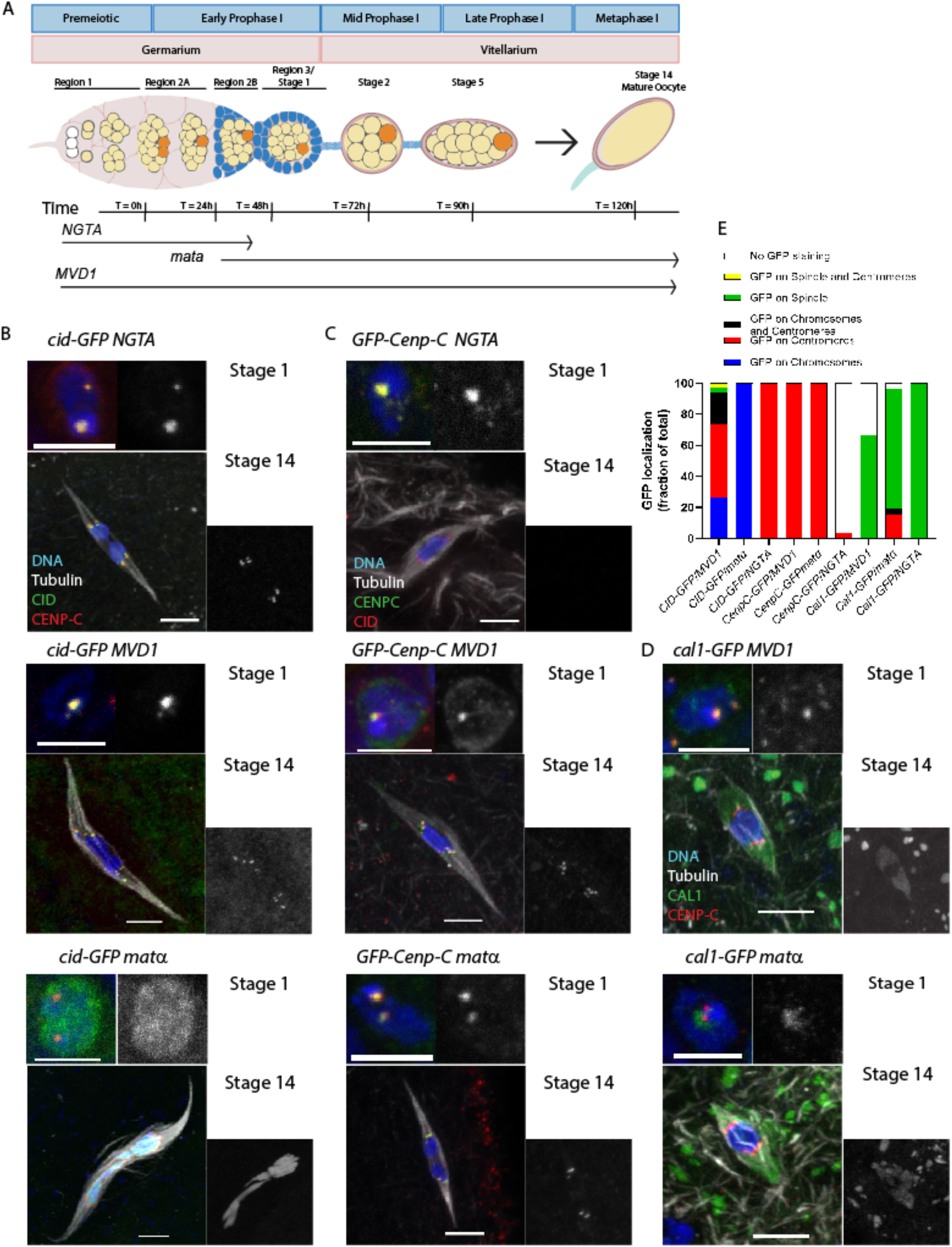
Loading of centromere proteins during meiotic prophase in oocytes. **A)** Oocytes are generated and enter meiotic prophase in the germarium. Region 1 contains the stem cell niche as well as the pre-meiotic cyst cells undergoing mitosis. Prophase I of meiosis begins in region 2A. Time points are relative to the onset of meiosis [33, 34]. Region 3 is also known as vitellarium stage 1, and at this stage, a single oocyte (orange) has been determined. *NGTA* begins expressing in the mitotic germline (region 1) and ends in region 3 or early in the vitellarium. *MVD1* expression begins earlier than *NGTA,* being expressed in all the mitotic and meiotic stages of the germline, including the stem cells. *Mata* begins expressing in region 2B or 3 and continues until stage 14, the metaphase I-arrested oocyte [36]. **B)** Localization of GFP-tagged CID (green) and centromeres detected using a CENP-C antibody (red). **C)** Localization of GFP-tagged CENP-C, with the centromeres detected using a CID antibody (red). **D)** Localization of GFP-tagged CAL1 (green) with the centromeres detected using a CENP-C antibody (red). In all images, the DNA is blue and the scale bars are 5 um. **E)** Summary of CID, CENP-C, and CAL1 localization in Stage 14 oocytes (n = 34, 18, 11, 12, 10, 30, 15, 26, 4).

To determine if CENP-C, CAL1, or CID can assemble on chromosomes during meiotic prophase, we used *UASP* regulated EGFP-tagged transgenes. With these transgenes, expression of *Cenp-C, cal1,* and *cid* was induced during female meiotic prophase using one of three GAL4-expressing transgenes (Figure 1A). We started with *P{GAL4::VP16-nos.UTR}CG6325^MVD1^* (referred to as *MVD1*) because it promotes expression of UAS transgenes through all stages of the germline, including the pre-meiotic cyst cells of the germline and continuing throughout all stages of oocyte development [35]. Crossing each transgene to *MVD1* resulted in CID, CAL1, and CENP-C localization in region 2a and throughout stages 1 to 5 of oogenesis (Figure 1B-D, Figure S 1).

As expected, CENP-C regulated by *MVD1* (referred to as *GFP-Cenp-C/MVD1* oocytes) was localized to meiotic centromeres in metaphase I of stage 14 oocytes. In contrast, surprising patterns were observed in stage 14 oocytes when CID or CAL1 were regulated by *MVD1* (Figure 1B, D). In *cid-GFP/MVD1* oocytes, CID was present as centromeric foci in about 50% of oocytes (Figure 1E). In other oocytes, however, CID was present on most of the meiotic chromatin. It was enriched in puncta and appeared to recruit CENP-C to these ectopic sites (Figure S 2). Thus, overexpression of CID in prophase can result in non-centromeric chromatin localization. In contrast, CAL1 was absent from the centromeres in *cal1-GFP/ MVD1* stage 14 oocytes, although it was frequently on the spindle (Figure 1D, Figure S 2). The spindle localization appears to be due to overexpression because it was not observed when CAL1 was regulated by its own promoter (Figure S 2). Thus, CAL1 appears to be unloaded from meiotic centromeres during prophase.

To test if early prophase loading was sufficient for later stages of meiosis, we used *P{GAL4-nos.NGT}A* (referred to as *NGTA*), which expresses in the germarium, including the mitotic divisions and early meiotic prophase (Figure S 3). Importantly for our study, *NGTA* expression ends in early prophase (stages 3-5). When regulated by *NGTA,* centromeric localization of CID, CAL1, and CENP-C was observed in region 2a and stages 1-5 (Figure 1B, C, Figure S 1). Similar results were observed with *HA-Cenp-C/ NGTA* oocytes (Figure S 4A). These results correspond to the *NGTA* expression pattern. In stage 14 oocytes, considerably after the zone of *NGTA* expression, CID was localized at the centromeres (Figure 1B). These data show that CID loaded during or prior to early meiotic prophase is maintained throughout prophase and into metaphase I of stage 14 oocytes. As with *MVD1,* CAL1 was not detected at centromeres in *cal1-GFP/ NGTA* stage 14 oocytes (Figure S 1), confirming that CAL1 is unloaded during meiotic prophase. While *GFP-Cenp-C/ NGTA* oocytes had centromeric CENP-C localization in stages 1-5 (see above), it was absent in stage 14 oocytes (Figure 1C). These results suggest that CENP-C is unloaded from the centromeres during meiotic prophase.

To test if centromere proteins load during meiotic prophase, we used *P{w[+mC]=matalpha4-GAL-VP16}V37* (referred to as *mata*), which begins expression in late prophase (region 2B/3) and continues through metaphase I in the stage 14 oocyte [36, 37]. In *cal1-GFP/ mata* oocytes, a weak GFP signal was frequently observed on the spindle, and rarely at the centromeres (Figure 1D, Figure S 2). This is consistent with the observations made in *cal1-GFP/ MVD1* oocytes that CAL1 is absent from metaphase I centromeres. CID induced by *mata* during late prophase I was present on most of the meiotic chromatin in 100% of the oocytes (Figure 1B). This was similar to the pattern observed in some of the *cid-GFP/ MVD1* oocytes; CID was enriched in some puncta, and also appeared to recruit CENP-C and SPC105R to the meiotic chromatin (Figure S 2). Ectopic CID affected chromatin organization, as individual bivalents could be discerned, unlike wildtype metaphase I oocytes in which the chromosomes are assembled in a single karyosome. These females were also sterile, suggesting that overexpression of CID during prophase appears to be deleterious, resulting in CID localizing to most of the chromatin, recruiting other centromere and kinetochore proteins, and causing changes in chromosome organization.

CENP-C localized only to the centromeres in *GFP-Cenp-C/ mata* oocytes (Figure 1D, Figure S 4A,B). In addition, heat shock was used to induce a pulse of expression during prophase and resulted in localization of CENP-C (Figure S 4C). These data indicate that CENP-C is recruited to centromeres during meiotic prophase. Thus, meiotic CENP-C dynamics are different than CID and CAL1. CID expression during or prior to early prophase is sufficient for meiosis I. In contrast, CENP-C expression later in prophase is required for its centromere localization in meiosis I.

### Centromeric CENP-C exchanges during prophase I

The results with *NGTA* and *mata* used to express *GFP-Cenp-C* suggest that centromeres unload and load CENP-C during meiotic prophase. To test the relationship between the unloading and loading of CENP-C, females were generated that expressed two forms of CENP-C (Figure 2A). *GFP-Cenp-C* was under the control of its own promoter while *HA-Cenp-C* was under the control of *mata*. With these transgenes we could measure if the HA-CENP-C induced in prophase would replace the constitutively expressed GFP-CENP-C. In addition, we knocked down *GFP-Cenp-C* expression in prophase by RNAi using shRNA *GL00409,* which targets a sequence within the 5’ UTR of *Cenp-C* and is not present in *HA-Cenp-C*. GFP-CENP-C was expected to be expressed at all stages of the germline but, due to containing the 5’ UTR, was sensitive to RNAi in meiotic prophase. In contrast, HA-CENP-C expression would begin in early prophase and be resistant to RNAi because it lacks the 5’UTR. This setup allowed us to determine whether unloading of centromeric CENP-C is dependent on a cytoplasmic pool of CENP-C.

**Figure 2:**
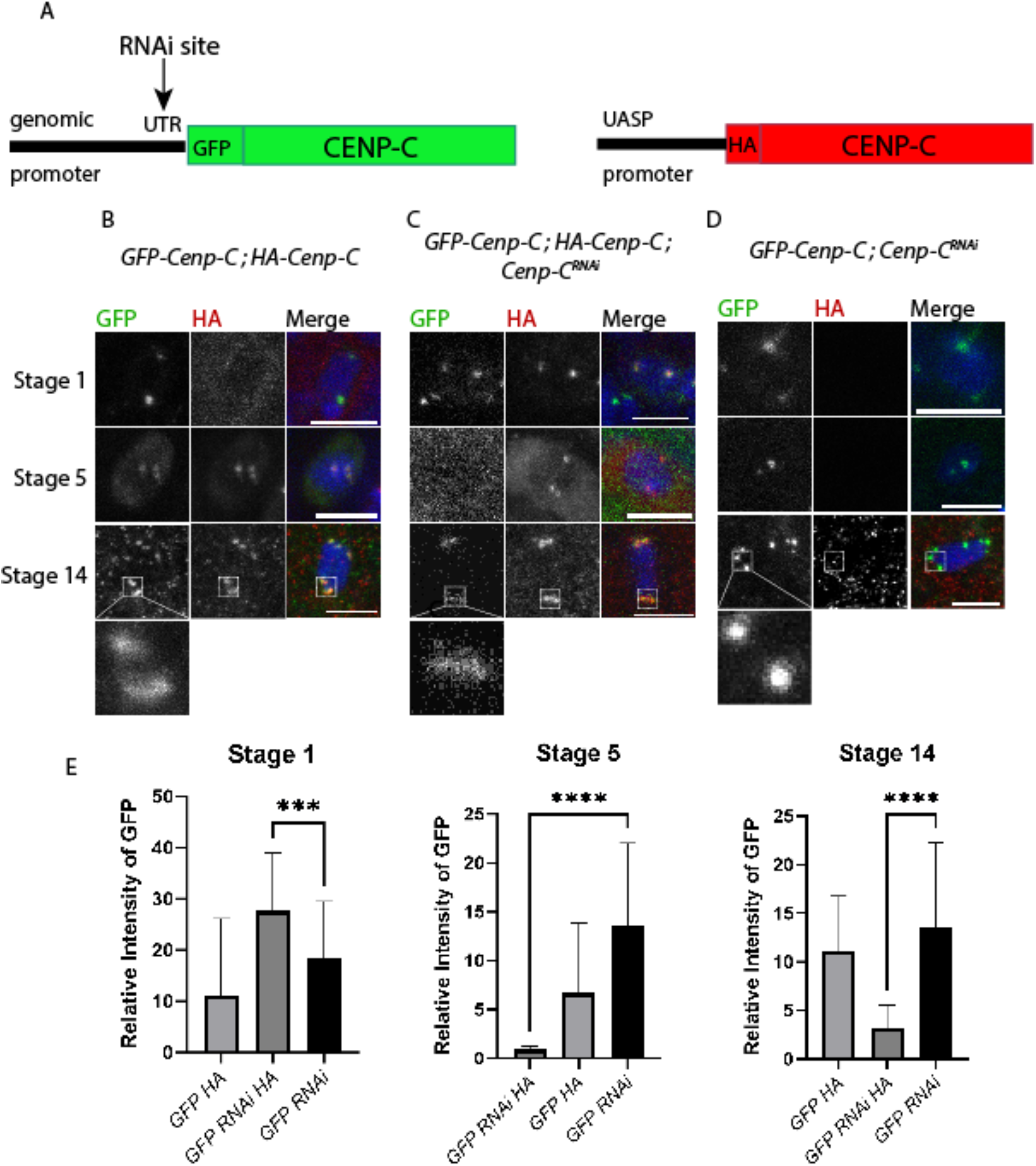
CENP-C is exchanged at the centromeres during meiotic prophase. Immunocytology was performed on oocytes ubiquitously expressing GFP-CENP-C (green), plus or minus *GL00409* (*Cenp-C^RNAi^*), and HA-CENP-C (red) using the *mata* promoter. All scale bars represent 5 um and DNA is blue. **A)** *GL00409* recognizes a sequence in the 5’ UTR, and, therefore, degrades *GFP-Cenp-C* but not *HA-Cenp-C*. Images show stages 1, 5 and 14 with (**B)** expression of GFP-CENP-C and HA-CENP-C, **(C)** expression of GFP-CENP-C, *Cenp-C^RNAi^*, and HA-CENP-C, and (**D)** expression of GFP-CENP-C with *Cenp-C^RNAi^*. **E)** The relative intensity of GFP-CENP-C was measured at each stage in each genotype (n = 70, 117, 28, 23; 12, 19, 104, 156; 50) with error bars showing standard deviation. For each genotype, the intensity between stage 1 and stage 14 was compared using an unpaired t-test (***p = 0.001; ****p<0.0001).

In *GFP-Cenp-C*, *HA-Cenp-C* (no RNAi) females, the GFP and HA variants were both maintained (Figure 2B, E). A severe reduction in stage 5 and stage 14 GFP-CENP-C levels was observed in *GFP-Cenp-C*, *HA-Cenp-C*, *GL00409* females (Figure 2C, E). In contrast, the decrease in GFP-CENP-C was less severe in the absence of HA-CENP-C expression (*GFP-Cenp-C*, *GL00409* females, Figure 2D, E). To explain these observations, we propose that the unloading of centromeric CENP-C depends on the availability of a replacement. In addition, while the GFP-CENP-C levels at the centromeres drops dramatically in the presence of HA-CENP-C and RNAi, a low level was still observed in stage 14 oocytes (Figure 2D). These results suggest that, although most CENP-C is exchanged during prophase, there is a small pool of CENP-C that is maintained throughout meiosis.

### CENP-C regulates centromere clustering, recombination, and MIS12 loading in early prophase I

These results show that there is a population of CENP-C loaded during meiotic prophase that is distinct from the population required for the maintenance of CID at the centromeres. To investigate *Cenp-*C function in early prophase, we knocked down its expression by RNAi using an shRNA (*GL00409*) with either the *NGTA* or *MVD1* promoters. Females expressing *GL00409* with *MVD1* (*GL00409*/*MVD1* oocytes) were sterile and agametic [38]. *MVD1*-driven expression in all premeiotic mitotic germ cells and a requirement for CENP-C during these mitotic divisions would explain the agametic phenotype. Thus, the remaining early prophase experiments were performed with females expressing *GL00409* with *NGTA* (*GL00409/NGTA* oocytes).

*GL00409/NGTA* females were fertile but displayed a high level of X-chromosome nondisjunction (7%) compared to controls (0.25%) (Table 1). A similar result was observed with a different shRNA targeting *Cenp-C* (*HMJ21500*). Because *NGTA* expression occurs only during early prophase, these results suggest CENP-C has an early meiotic function required for chromosome segregation. To investigate the mechanisms of nondisjunction caused by loss of CENP-C, we characterized processes that occur early in meiotic prophase such as centromere assembly, meiotic recombination, centromere pairing, sister chromatid cohesion, and synaptonemal complex (SC) formation.

**Table 1:** Fertility, nondisjunction and crossing over in *Cenp-C* RNAi females.

|  | Fertility | Nondisjunction (NDJ) |  | Recombination (% of control) |  |  |  |
| --- | --- | --- | --- | --- | --- | --- | --- |
|  | Offspring per female<br>(# of females) | Homologous Chromosome NDJ (# of offspring) | Sister Chromatid NDJ (# of offspring) | *st-cu*<br>(*m.u.*) | *cu-e*<br>(*m.u.*) | *e-ca*<br>(*m.u.*) | # of progeny |
| Control | 24.3 (105) | 0.25% (1590) | 0.5% (2696) | 4.4 | 2.2 | 29.4 | 2853 |
| *GL00409 / NGTA* | 10.0 (240) | 7.0% (1658) | 2.6% (1164) | 9.4<br>(226%) | 21.1<br>(101%) | 22.9<br>(81%) | 1668 |
| *HMJ21500 / NGTA* | 3.3 (133) | 7.6 (1213) | ND | ND | ND | ND | ND |
| *GL00409 / MVD1* | 0.27 (105) | 0% (54) | -<sup>a</sup> | - | - | - | - |

<sup>a</sup> not enough progeny to accurately measure nondisjunction or crossing over

The intensities of CENP-C and CID at the centromeres were moderately decreased in early prophase nuclei of *GL00409*/*NGTA* oocytes (Figure 3A - C). The mild reduction of both centromere proteins suggests that the knockdown of CENP-C only had a mild effect on centromere maintenance. For another measure of CENP-C function in prophase, we examined the localization of MIS12. In *Drosophila* mitotic cells, CENP-C promotes kinetochore assembly by recruiting MIS12 [23–25]. MIS12-GFP localized to centromeres in control oocytes, starting in region 2a of the germarium (Figure 3D). The intensity of centromeric MIS12-GFP was significantly reduced in *GL00409/NGTA* females (Figure 3E) indicating that CENP-C promotes recruitment of MIS12 during meiotic prophase. These results show that MIS12 localizes to the centromeres in meiotic prophase and depends on CENP-C.

**Figure 3.**
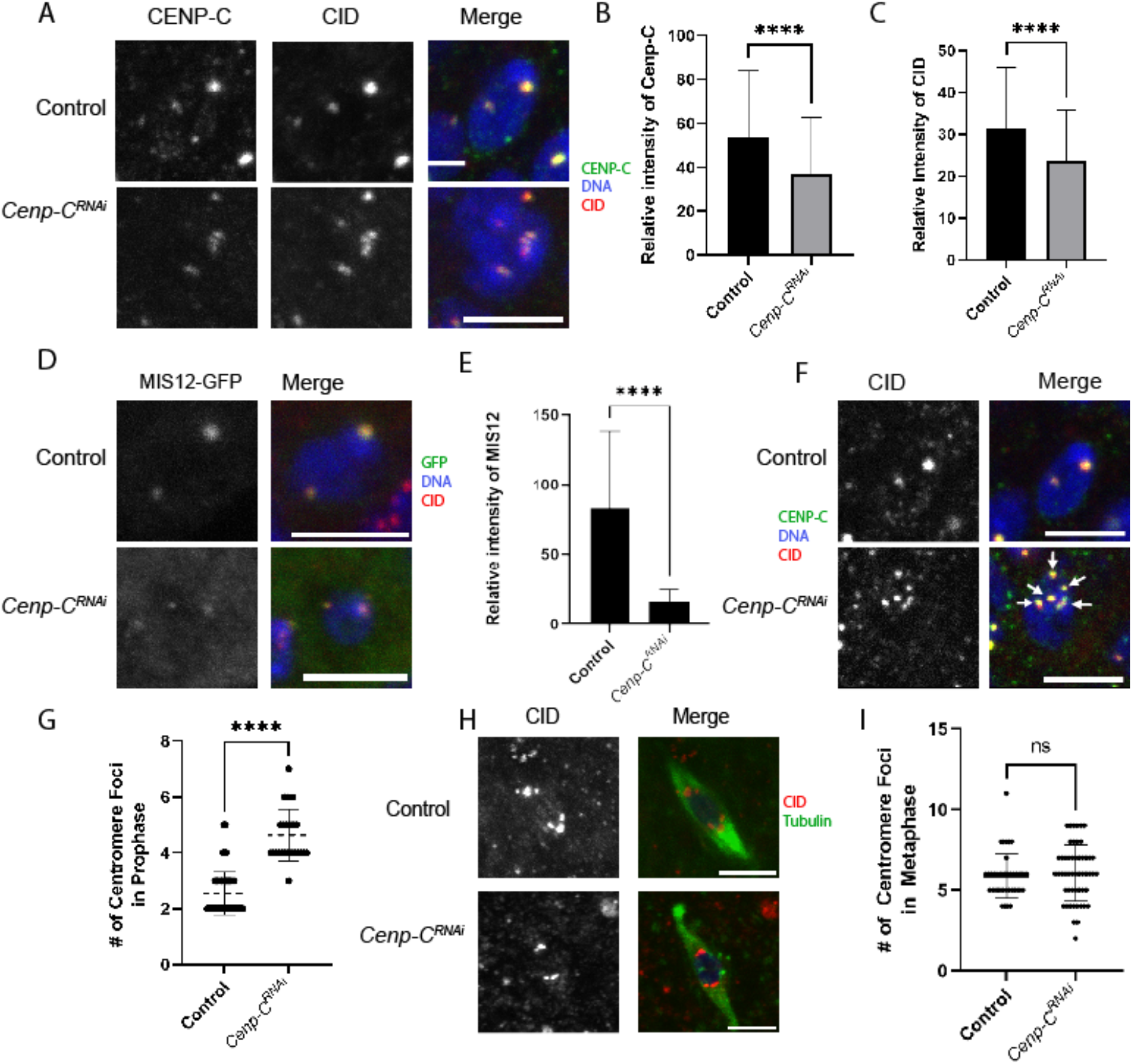
Early Prophase I defects when CENP-C is depleted using NGTA. Immunocytology was performed on *GL00409/NGTA* oocytes with CID (red), DNA (blue), and scale bars represent 5 um. **A)** Region 2 oocytes showing CENP-C in green. **B-C)** The intensities of CENP-C and CID in region 2 were measured in *Cenp-C* RNAi and control oocytes (For B, n = 104 and 88; for C, n = 104 and 87). **D)** Region 2 oocytes with MIS12-GFP in green. **E)** MIS12 intensity was measured in region 2 (n = 13 and 30). **F)** CID foci in *Cenp-C* RNAi region 3 oocytes, with CENP-C in green. **G)** Number of centromere foci was measured based on CID foci in contact with the DNA (n = 84 and 27). **H)** Tubulin (green) and CID foci (red) in *Cenp-C* RNAi metaphase I oocytes. **I)** There was no increase in CID foci observed in stage 14 oocytes (n = 40 and 52). Error bars represent standard deviation from the mean. ****p<0.0001

Increased frequencies of nondisjunction can be associated with loss of SC and/or a reduction in crossing over (CO) [39]. SC assembly in *GL00409/NGTA* females was normal, indicated by the thread-like appearance of the transverse filament protein C(3)G [40] (Figure S 5). We measured the frequency of crossing over in *GL00409/NGTA* females and observed an increase in centromeric crossovers, but overall, crossing over was not decreased (Table 1). When the centromeres were genetically marked (see Methods, Table 1), we observed an increase in sister chromatid nondisjunction. Thus, meiotic nondisjunction may occur because of a cohesion defect, which could be caused by the increase in proximal crossovers [41].

A cohesion defect can be observed as an increase in the number of centromere foci [42, 43]. Indeed, there was an increased number of CID foci in the germaria of *GL00409/NGTA* ovaries, indicating a separation of centromeres (Figure 3F, G). We tested if the cohesion defect persisted, and therefore would result in increased CID foci in metaphase I oocytes. The variability in CID foci number was greater in the experimental group, but the mean was not significantly different from the control (Figure 3H, I).

Because nonhomologous centromeres normally cluster in early prophase, we used chromosome-specific fluorescence in situ hybridization (FISH) probes for the pericentromeric regions of the X and 2^nd^ chromosomes to examine pairing and clustering of the centromeres (Figure 4A). Clustering defects were defined as nuclei containing 3 or more clearly separated probe foci (of any color) whereas control oocytes had 1-2 foci. As with the experiment to detect CID foci, the FISH foci failed to cluster (Figure 4B). Pairing defects were defined as greater than one FISH focus per probe. While homologous centromeres were efficiently paired in wild-type germarium, pairing was defective in *GL00409/NGTA* oocytes (Figure 4C). These results suggest that there are defects in both centromere pairing and clustering, consistent with a defect in sister centromere cohesion.

**Figure 4:**
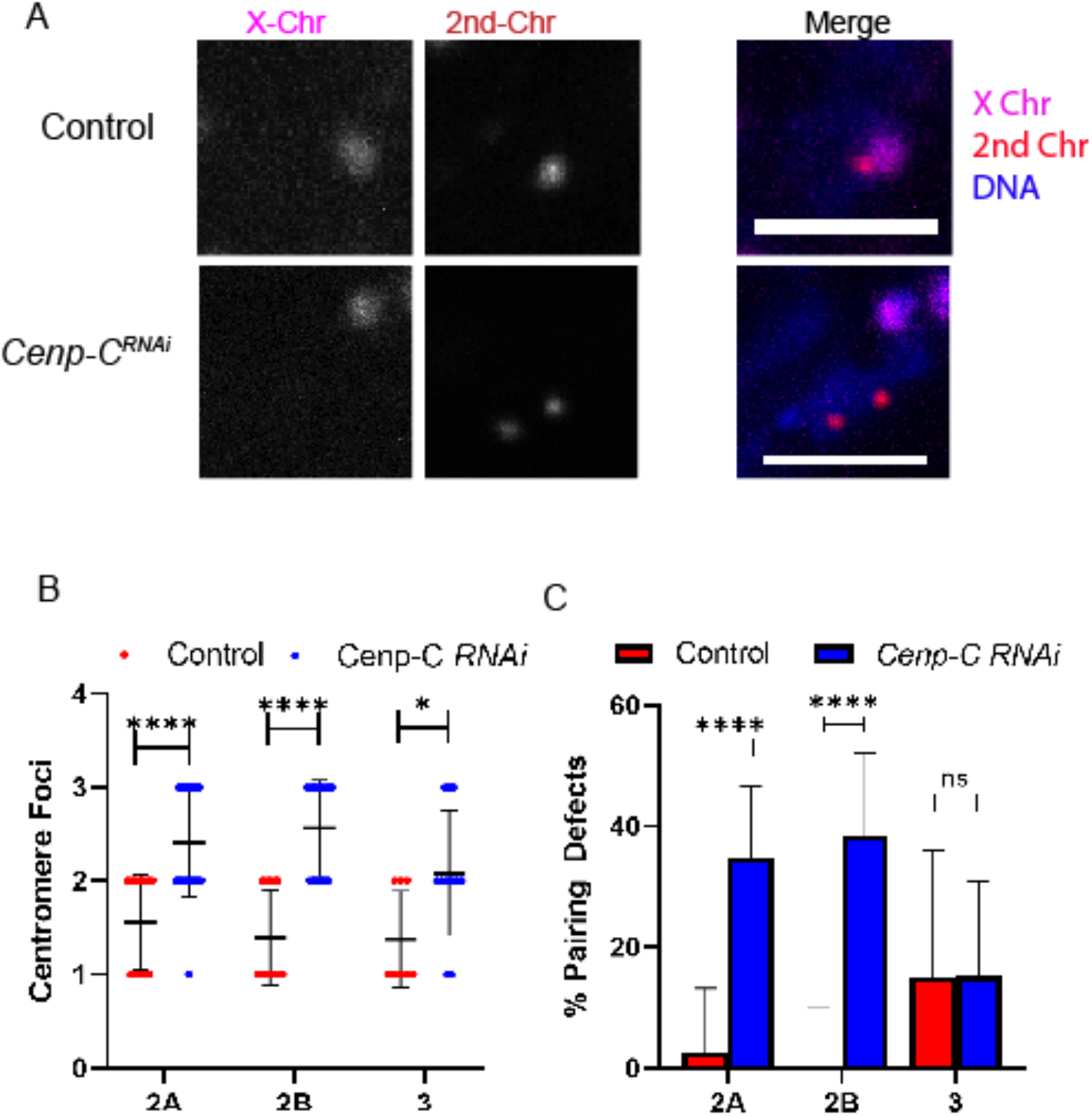
CENP-C is required for centromere pairing and clustering. **A)** FISH probes were used to detect the pericentromeric regions of chromosome 2 (red), and X (magenta) in the germarium of *GL00409/NGTA* or control oocytes. DNA is in blue. **B)** Clustering defects were defined as nuclei with greater than 2 of any centromere foci (control n = 16, 20, 14; RNAi n = 42, 38, 18). **C)** Pairing defects were defined as oocytes with greater than one focus for a chromosome in a nucleus. Scale bars represent 5 µm and error bars show standard deviation. * = 0.0196>p>0.0238, ****p<0.0001.

### Prophase CENP-C plays a critical role in maintenance of centromeres and kinetochores

To test the function of CENP-C loaded during late prophase, we used *mata* to express *Cenp-C* shRNA (*GL00409/mata*). The *GL00409/mata* females were sterile, although this could be due to defects in early embryo mitosis rather than meiosis. To investigate the effects on meiosis I, stage 14 *GL00409/mata* oocytes were analyzed for meiotic defects. The intensity of CENP-C at the centromeres was reduced to approximately 25% of the intensity in control oocytes (Figure 5A, B). The intensity of CID at the centromeres was mildly reduced in *GL00409/mata* oocytes to approximately 80% of control levels (Figure 5A, C). These results suggest that CENP-C might play a role in the stabilization of CID late in meiotic prophase or in metaphase I.

**Figure 5:**
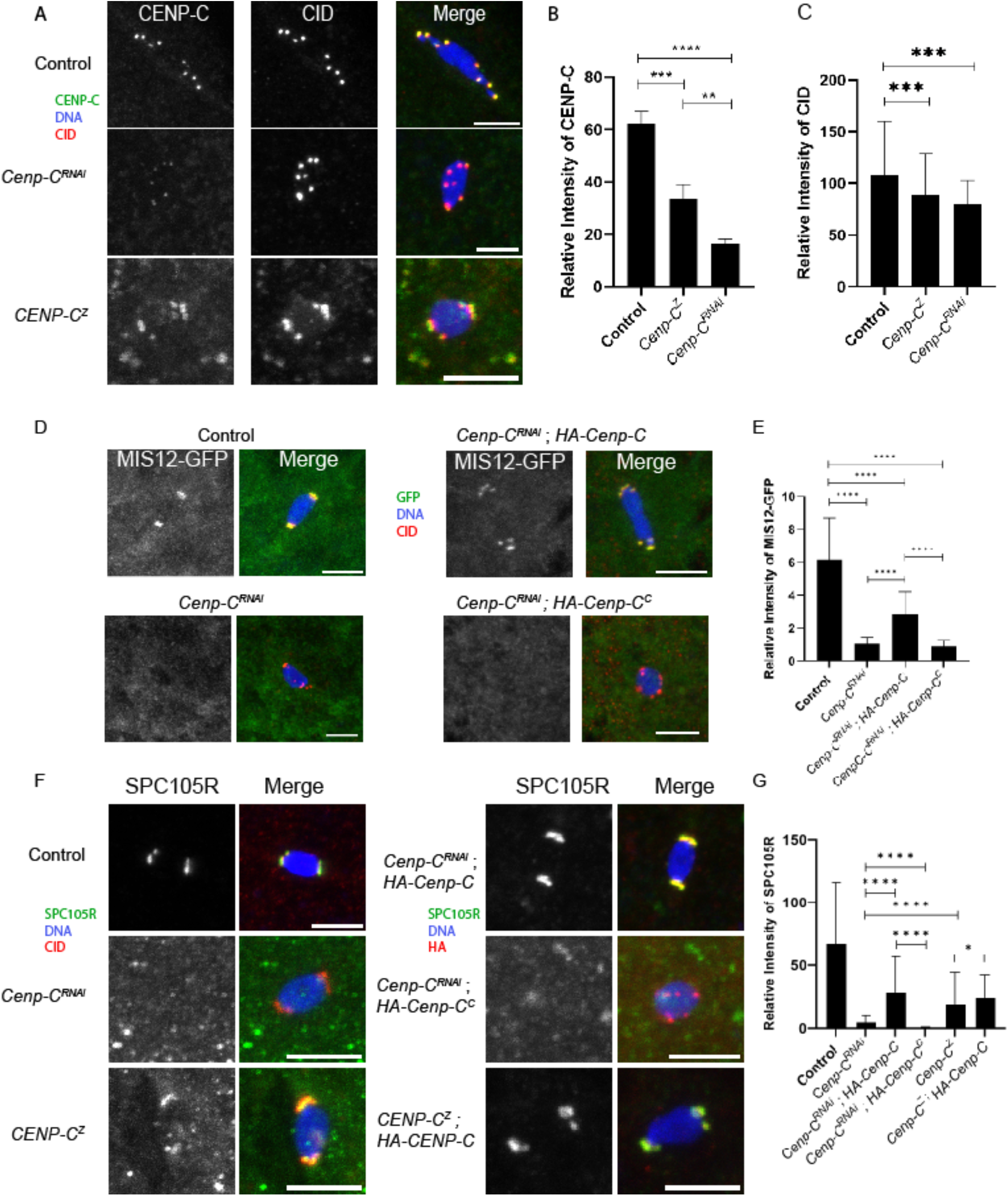
CENP-C is required for kinetochore assembly and chromosome segregation. Immunocytology was performed on stage 14 oocytes expressing the *GL00409* with *mata* (referred to as *Cenp-C^RNAi^*) or heterozygous for *Cenp-C* mutations (*Cenp-C^Z3-4375^/Cenp-C^IR35^* referred to as “*Cenp-C^Z^*”). HA-tagged *Cenp-C* transgenes were coexpressed using the *mata* promoter in RNAi experiments or *osk-Gal4* in *Cenp-C^Z^* experiments. DNA is in blue and scale bars represent 5 um in all images. **A)** Oocytes with CENP-C (green) and CID (red). **B)** The intensity of CENP-C was measured relative to background (n = 55, 67, 54). **C)** The intensity of CID was measured relative to background (n = 186, 221, 54). **D)** Oocytes expressing MIS12-GFP (green) and CID (red). **E)** Mis12-GFP localization was measured relative to the background (n = 57, 41, 47, 67). **F)** Oocytes with SPC105R (green) and CID (red). **G)** Intensity of SPC105R was measured relative to background (n = 108, 136, 115, 132, 245, 150). *p = 0.0379, **p = 0.0086, *** = 0.0002>p>0.0001, ****p<0.0001

To determine if the loss of CENP-C affects kinetochore assembly, we quantified the intensity of MIS12 and SPC105R in mature oocytes. Consistent with the reduction in CENP-C levels, we observed approximately 85% reduction in MIS12 levels and a 90% reduction in SPC105R in *GL00409/mata* oocytes (Figure 5D-G). Consistent with the loss of SPC105R recruitment, we observed an increase in the distance between centromeres and microtubules (Figure S 6), which has previously been observed when SPC105R is depleted [44]. These results suggest CENP-C recruits MIS-12, which then recruits SPC105R, although we can’t rule out CENP-C recruits SPC105R directly. We previously showed that loss of SPC105R was associated with loss of sister centromere cohesion [45]. However, we did not observe this phenotype in *GL00409/mata* oocytes (Figure S 6), suggesting sufficient SPC105R remains to protect cohesion.

To determine whether the loss of CENP-C in oocytes affected chromosome segregation at meiosis I, we assayed for bi-orientation of homologous chromosomes using FISH. We observed a dramatic increase in bi-orientation errors (primarily mono-orientation) in *GL00409/mata* oocytes (Figure 6A,B), and a decrease in the distance between homologous centromeres, compared to the control (Figure 6C). These spindle attachment errors could indicate that CENP-C plays a direct role in error correction, but more likely has an indirect effect due to the function of CENP-C in kinetochore assembly.

**Figure 6.**
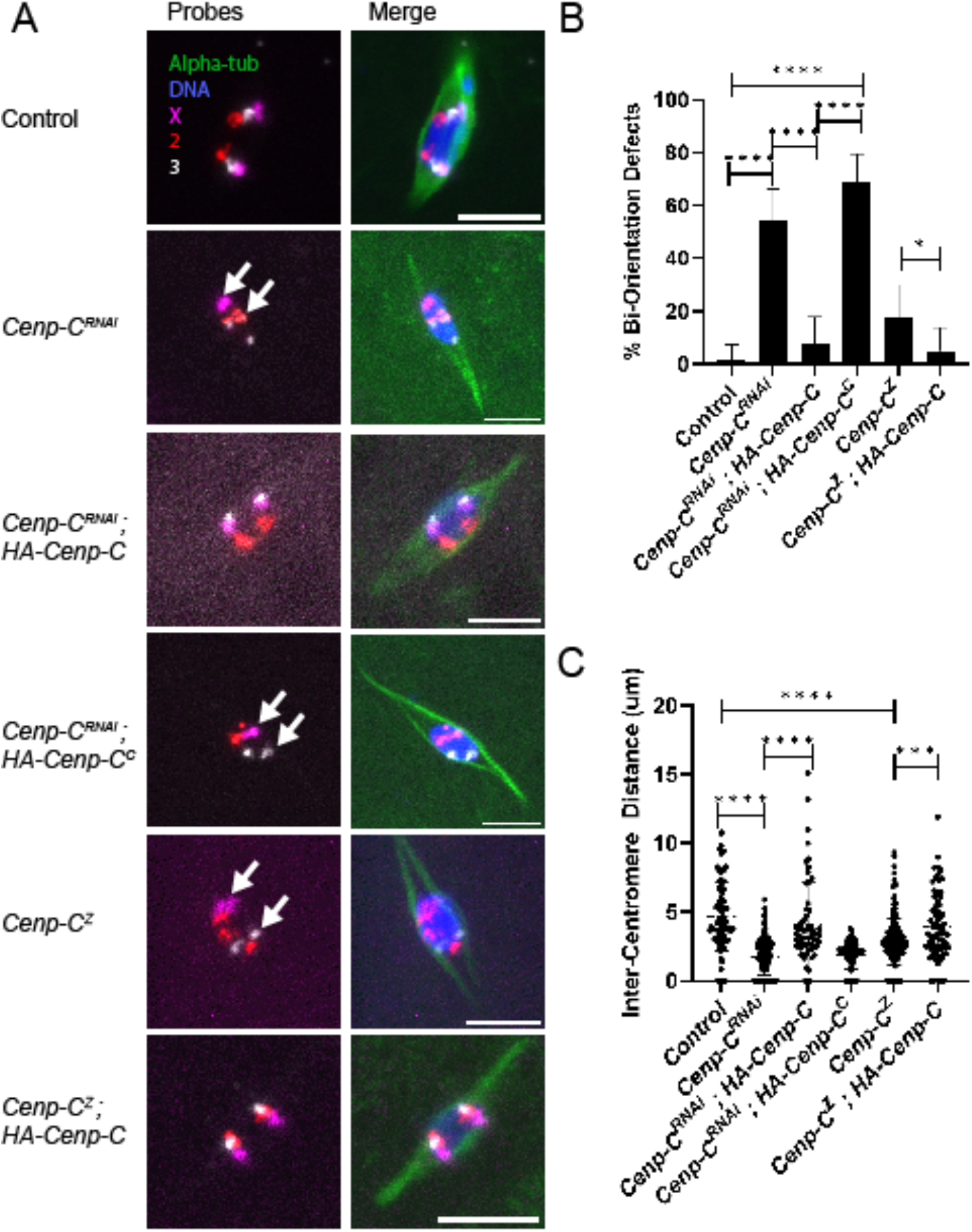
Prophase-loaded CENP-C is required for bi-orientation in meiosis I. **A)** FISH probes were used to detect the pericentromeric regions of chromosome 2 (red), 3 (grey), and X (magenta). Stage 14 oocytes were *Cenp-C^Z3-4375^/Cenp-C^IR35^* (*Cenp-C^Z^*) or *GL00409/mata* (*Cenp-C^RNAi^*). In some cases, *HA-Cenp-C* was expressed using *mata* in *Cenp-C^RNAi^* experiments or *oskGal4* in *Cenp-C^Z^* experiments. **B)** Bi-orientation defects were defined as two foci of the same probe that were on the same side of the spindle midzone or only a single focus, indicating mono-orientation (n = 75, 59, 53, 58, 56, 60). **C)** The distance between the center mass of two foci of the same probe, which measures the separation of homologous chromosomes towards opposite poles (n = 90, 156, 60, 60, 192, 72). All scale bars represent 5 um and error bars represent standard deviation from the mean. * = 0.0473>p>0.0387, **p = 0.0024, *** = 0.0002 >p> 0.001, ****p<0.001.

### Rescue of the CENP-C phenotype by late prophase expression of a wild-type allele

Prophase-expressed CENP-C can load onto the centromeres and the defects in *GL00409*/*mata* females suggest that prophase CENP-C expression is required for meiosis and fertility. To directly test if prophase-expressed CENP-C is functional in meiosis, we tested if the *HA-Cenp-C* transgene could rescue loss of endogenous *Cenp-C*. The *HA-Cenp-C* transgene lacks the 5’UTR that includes the target sequence for *GL00409,* and therefore, is resistant to knockdown by RNAi. When *HA-Cenp-C* was co-expressed with *GL00409* using *mata* in oocytes, we observed localization of HA-CENP-C to the centromeres and restoration of MIS12 and SPC105R localization (Figure 5D-G). In addition, the biorientation defects in *GL00409/mata* oocytes were rescued by expression of HA-CENP-C (Figure 6). Thus, prophase expression of a CENP-C transgene rescued the defects observed in RNAi oocytes.

Despite the rescue of the meiotic RNAi phenotypes, however, expression of HA-CENP-C using *mata* caused sterility (Table 2). These results suggest overexpression of CENP-C leads to loss of fertility. The females expressing HA-CENP-C had normal oocytes at metaphase I, making it likely that the defect causing sterility is after meiosis I, possibly during embryonic mitosis. To circumvent the sterility from *mata-*induced expression of *Cenp-C,* we used *P{w[+mC]=osk-GAL4::VP16}A11* (referred to as *oskGal4*). The expression pattern of *oskGal4* is similar to *matα* but at lower levels [46]. Using *oskGal4* to induce *UASP-*regulated *Cenp-C*, either GFP-tagged or HA-tagged, resulted in fertile females (Table 2). We then used *oskGal4* to determine if prophase expression of CENP-C could rescue the phenotype of a hypomorphic allele, *Cenp-C^Z3-4375^* [47].

**Table 2:**
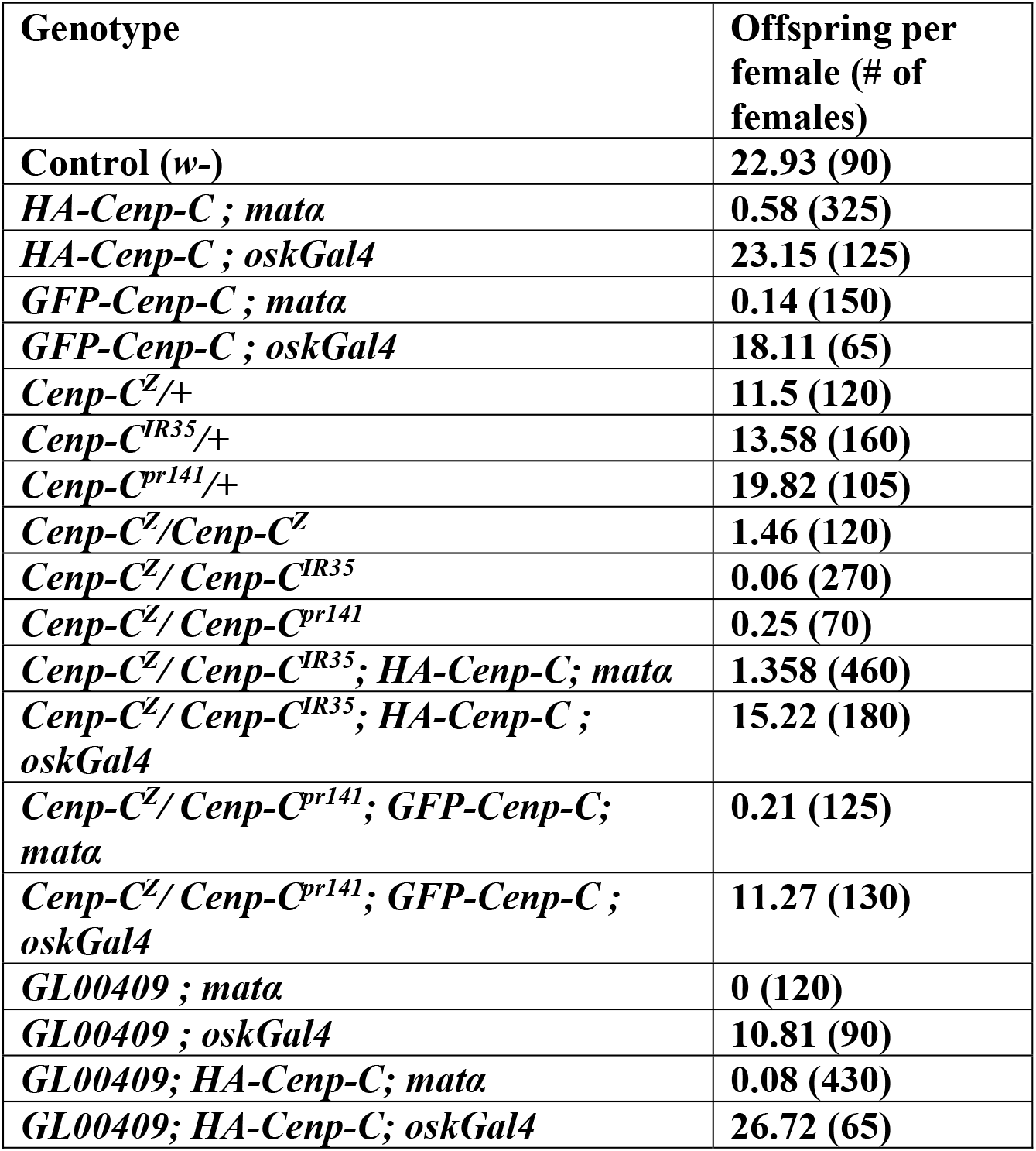
Fertility in *Cenp-C* RNAi, mutant and transgenic females.

When *Cenp-C^Z3-4375^* was heterozygous to a null allele (*Cenp-C^IR35^*), the females were sterile and centromeric CENP-C levels in oocytes were reduced, though not to the same levels as with *GL00409* (Figure 5A, B). Bi-orientation errors were also elevated in *Cenp-C^Z3-4375^/Cenp-C^IR35^* mutant oocytes, though not to the same extent as the shRNA-expressing oocytes (Figure 6). Expression of HA-CENP-C or GFP-CENP-C using *oskGal4* was found to rescue the sterility of *Cenp-C^Z3-4375^/Cenp-C^IR35^* mutant females (Table 2). In addition, SPC105R localization was restored (Figure 5F-G) and the biorientation defects in *Cenp-C^Z3-4375^/Cenp-C^IR35^* mutant females was rescued (Figure 6). These results show that late prophase expression of CENP-C contributes to kinetochore function in meiosis I.

### Regulation of CENP-C loading and centromere integrity

The RNAi resistance of the *HA-Cenp-C* transgene can be used to express mutant variants in the absence of wild-type CENP-C. Previous studies have shown that the C-terminal domain of *Drosophila* CENP-C interacts with the centromere components CID and CAL1 [17, 23]. To determine what domains of CENP-C are required for prophase loading in oocytes, we generated two *Cenp-C* transgenes expressing either the N-terminal (aa 1-788, *Cenp-C^N^*) or C-terminal (789–1411, *Cenp-C^C^*) domain of CENP-C. The N-terminal domain was not detected by fluorescence microscopy, suggesting it is either unstable or does not localize. In contrast, CENP-C^C^ was loaded to the centromeres during prophase (Figure S 4, Figure S 6), but failed to recruit SPC105R or MIS12 (Figure 5D-G). The CENP-C*^C^* protein may have a dominant negative effect because there was a significant reduction of SPC105R localization in *GL00409; Cenp-C^C^*oocytes compared to *GL00409.* These results support the conclusion that the C-terminal domain of CENP-C is required for centromere localization, while the N-terminal domain recruits MIS12 and the rest of the outer kinetochore.

Two pathways have been proposed to load CENP-C via the C-terminal domain. The first occurs during anaphase or G1, is required to maintain centromere identity [8, 48], and depends on CAL1 [10]. The second occurs during prophase, may be required to build kinetochores, and depends on CDK1 in mitotic cells [49]. To examine the role of CAL1 in prophase loading, we measured CENP-C loading in *cal1* RNAi oocytes (*GL01832*). Similarly, the same experiment was done using an shRNA (*HMS01531*) to knock down *Cdk1* mRNA. In both experiments, HA-CENP-C was observed in stage 5 oocytes, suggesting neither CAL1 nor CDK1 are required for prophase loading of CENP-C (Figure S 7). We did not examine stage 14 oocytes because, as described above, CAL1 is not present, and *Cdk1* RNAi oocytes fail to enter prometaphase.

The *Cenp-C^Z3.4375^* mutation is in the C-terminal domain and is present at the centromeres in decreased amounts (Figure 5A, B). We also observed a second phenotype suggesting a loss of kinetochore integrity. In some *Cenp-C^Z3.4375^* mutant oocytes, foci of CENP-C protein were detached from the main chromosome mass and moved towards the poles (Figure 7A, B). This suggests that one of the defects in *Cenp-C^Z3.4375^* mutant oocytes is a failure to maintain attachment of the kinetochore to the underlying centromeric chromatin. Surprisingly, this phenotype was observed in the presence of a wild-type transgene (Figure 7C). Either the kinetochore phenotype is dominant, or it is due to an early prophase function, and that expression of the transgene, which is in late prophase, was too late to rescue the phenotype.

**Figure 7:**
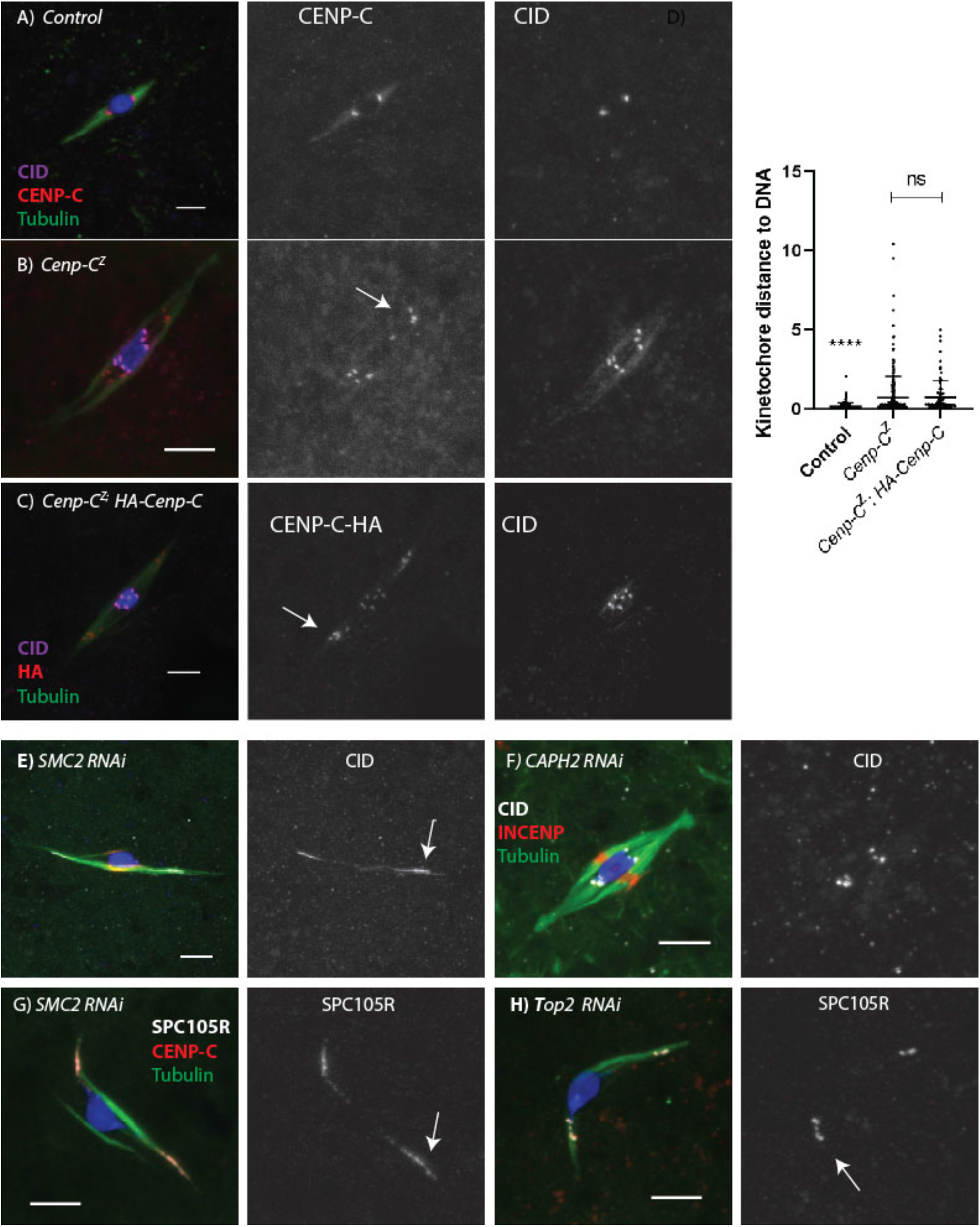
Kinetochore detachment in *Cenp-C^Z4.4375^* mutant oocytes. A-C) Kinetochore detachment in oocytes, with CENP-C in red, CID in magenta, microtubules in green, and DNA in blue. CENP-C foci have separated from the chromosomes (examples shown with arrows), while the CID remain**s** chromatin associated. Shown are (A) wild-type control, (B) *Cenp-C^Z3.4375 /^ Cenp-C^IR35^* (labelled *Cenp-C^Z^*), and (C) *HA-Cenp-C* transgene co-expressed in *Cenp-C^Z3.4375/^ Cenp-C **^pr141^*** mutant oocytes. D) The distance between the center of the kinetochore focus and the nearest surface of the DNA was measured. All scale bars represent 5 um and error bars represent standard deviation from the mean. (Control n=164 centromeres, *Cenp-C^Z3.4375/^ Cenp-C^IR35^* n=284, *Cenp-C^Z3.4375/^ Cenp-C^IR35^; HA-CENP-C* n=96). ****p<0.0001. D-E) Requirement of Condensin for kinetochore integrity. *CAPH2* or *SMC2* RNAi oocytes with CID (white), INCENP (red), tubulin (green), and DNA (blue). F-G) *SMC2* or *Top2* RNAi oocytes with SPC105R (white), CENP-C (red), tubulin (green), and DNA (blue). Scale bars represent 5 um.

The kinetochore detachment phenotype is reminiscent of results from budding yeast indicating condensins have a role in maintaining the cohesive structure of the centromeres [50, 51]. To compare the two phenotypes, we examined centromere and kinetochore protein localization in oocytes depleted of Condensin subunits. Depletion of SMC2 results in severe loss of centromere integrity (20/22 oocytes, Figure 7E). In contrast to *Cenp-C^Z3.4375^* mutant oocytes, CID was stretched towards the poles in addition to CENP-C. A similar, although less severe, phenotype was observed in *gluon* (SMC1) RNAi oocytes (5/11 oocytes), which could be due to a weaker knockdown. RNAi against subunits specific for Condensin I (Barren/Cap-H, 0/11) or Condensin II (CapH2, 0/6) did not have this phenotype, although we can’t rule out the possibility that the absence of a phenotype was due to insufficient mRNA knockdown (Figure 7F). SPC105R was also stretched towards the poles (Figure 7G), and a similar phenotype was observed using *Top2* RNAi (7/7, Figure 7H)), as observed previously [52]. These results show the importance of two mechanisms that resist the pulling forces on the kinetochores and keep the centromere and kinetochore together: Condensin /Top II maintains the compact centromeric chromatin while the C-terminal domain of CENP-C maintains the connection between centromeric chromatin and the inner kinetochore.

## Discussion

### Prophase dynamics of centromere proteins in oocytes

In *Drosophila*, centromere maintenance involves three proteins, CID, CAL1, and CENP-C, and occurs during late M or G1 (see Introduction). We show here that CENP-C must be loaded during meiotic prophase for centromere clustering, cohesion, and normal kinetochore assembly. We observed increased CENP-C unloading when there was a cytoplasmic source of CENP-C for replacement. This indicates that CENP-C dynamics involves an exchange mechanism, and without new sources of CENP-C, the dynamics are reduced. In contrast to CENP-C, expression of CID in premeiotic cells or early prophase was sufficient for robust localization in metaphase I. Surprisingly, CAL1 is unloaded during prophase and absent in metaphase I, which is in contrast to observations in mitotic cells [7, 16]. Compared to CID or CAL1, CENP-C is unique in requiring prophase loading for its functions in meiosis.

The stable property of CID is consistent with studies on CENP-A in mouse oocytes [28]. It has been reported, based on changes in fluorescence intensity, that CID is incorporated during meiotic prophase of *Drosophila* females [48]. These results do not conflict with ours, as we did observe small decreases in CID levels when CENP-C was depleted. However, the significance of this small amount of CID prophase loading is not known. In contrast, overexpression of CID during meiotic prophase resulted in ectopic localization to most of the chromatin, recruitment of kinetochore proteins like CENP-C and SPC105R, and sterility. Similar observations of ectopic localization have been made when CID is overexpressed in somatic cells [53–55]. Thus, it may be important to limit CID expression during prophase to avoid localization to non-centromeric locations.

The C-terminal domain of CENP-C had similar dynamic properties during prophase as the full-length protein. This part of the protein also contains the elements required to interact with CID and CAL1 for centromere maintenance [17, 18, 25]. Thus, CENP-C prophase loading may involve recruitment to existing sites containing CID. Some evidence in mammals, however, suggests CENP-C has both CENP-A-dependent and independent localization mechanisms, although in most of these studies, loading during G2 is not specifically addressed [56]. CDK1 has been implicated in CENP-C dynamics in chicken and human cells [49], although we did not see an effect on CENP-C loading when CDK1 was depleted in *Drosophila* oocytes. Further work is need to understand what regulates CENP-C dynamics during prophase.

### Early prophase function: centromere clustering and cohesion

When CENP-C was depleted from early prophase oocytes, we observed an increased number of centromere foci. Our results suggest the increased CID foci phenotype could be due to a loss of centromeric cohesion because we also observed an increase in sister chromatid nondisjunction. These observations are consistent with and extend prior studies on *Cenp-C* [47]. These authors also found that *Cenp-C^Z3-4375^* mutant females had defects in centromere clustering in pachytene oocytes (region 3 of the germarium). The centromere clustering and pairing phenotypes are similar to a hypomorphic allele of replication protein *mcm5* that was associated with loss of SMC1 at the centromeres [57]. Thus, we propose that CENP-C is required to recruit or maintain cohesion. A recruitment function of CENP-C could occur when cohesion is established in S-phase [42]. Alternatively, our results show striking similarities to the known roles of mammalian and yeast CENP-C in recruiting Moa1 or Meikin [58, 59], which are protectors of centromeric cohesion and particularly important for meiosis I.

### Late prophase function: kinetochore assembly

Depletion of prophase CENP-C resulted in phenotypes associated with loss of the kinetochore. This includes loss of MIS12 and SPC105R when CENP-C was depleted in prophase. These results show that CENP-C is required for most or all kinetochore assembly, consistent with studies in *Drosophila* mitotic cells [19, 24, 25]. These results also show that unlike CID/CENP-A, CENP-C loaded prior to prophase, that is required for centromere maintenance, is not sufficient for kinetochore assembly. Why CENP-C is loaded during prophase instead of prometaphase I like most other kinetochore proteins is not known. Studies in vertebrate cells have shown that CENP-C is dynamic during interphase but stable in metaphase [32, 60]. In addition, CENP-C turnover is much slower compared to microtubules and kinetochore proteins [61]. Therefore, it may be necessary to assemble CENP-C at the centromeres during prophase for two reasons: it is not stably maintained like CID and its dynamics are much slower than other spindle proteins. Prophase loading may also provide a stable and defined platform for kinetochore assembly at the beginning of meiotic prometaphase in oocytes.

The C-terminal domain of CENP-C was recruited to the centromeres but failed to assemble a kinetochore. Thus, the N-terminal domain of CENP-C appears to interact with MIS12 and recruit the rest of the kinetochore, consistent with previous studies in *Drosophila* [23, 27]. The *Cencp-C^Z3-4375^* mutation, which causes a substitution of amino acid 1115 [47], apparently weakens the interaction between CENP-C and the centromere. This is likely the cause of the detached kinetochore phenotype in this mutant, which indicates a failure to maintain kinetochore integrity when experiencing the force of microtubule pulling. We have also shown that Condensin, along with Topoisomerase II [52], is required to maintain the integrity of CENP-C and other kinetochore proteins on the chromosomes. Previous work in yeast has suggested that condensins have a role in maintaining the cohesive structure of the centromeres in metaphase [50, 51].

### Implications of CENP-C prophase loading

CENP-A/CID is remarkably stable [62, 63], not needing to be loaded during the extended prophase in oocytes. Our overexpression of CID results suggests that preventing CID from loading in prophase may be necessary to maintain a single centromere on each chromosome. In contrast, CENP-C must be loaded during prophase. Why CENP-C is not like CENP-A/CID and must be loaded during prophase is not known. It is possible that more CENP-C is needed for kinetochore assembly than centromere maintenance. It may simply be that, for unknown reasons, CENP-C is not as stable as CENP-A, and therefore requires replenishment to maintain its levels. Regardless of the reason, the requirement for CENP-C prophase loading could affect the health of aging oocytes. If loading of CENP-C is compromised in oocytes that have spent a longer time in meiotic prophase, our results show that sister chromatid cohesion, kinetochore assembly, and meiotic chromosomes segregation could be defective, leading to higher levels of aneuploidy.

## Acknowledgements

We thank Marina Druzhinina for technical assistance, Christian Lehner for providing Drosophila stocks and antibodies. We also thank Sarah Radford for early work on this project. We thank the Bloomington *Drosophila* Stock Center (NIH P40OD018537) and the TRiP project at Harvard Medical School for providing fly stocks used in this study. J.F was supported by a NIH IRACDA post-doctoral Fellowship. This work was supported by NIH grant GM101955 to K.S.M.

## Materials and Methods

### *Drosophila* strains and genetics

*Drosophila* crosses and stocks were kept at 25°C on standard medium. Fly stocks, including lines expressing a short hairpin RNA (shRNA), were obtained from the Bloomington Stock Center or the Transgenic RNAi Project at Harvard Medical School [TRiP, Boston, MA, USA, <u>flyrnai.org</u>]. Information about the genetic loci can be found on FlyBase [flybase.org]. *Drosophila* were obtained from the Bloomington Stock Center. Expression of the shRNA was controlled by the Upstream Activating Sequence (UAS), which is activated by expression of GAL4 under the control of a tissue-specific enhancer [1]. Five *Gal4* strains were used in this study, four of which were germline-specific: *P{matalpha4-GAL-VP16}V37* and *P{w[+mC]=osk-GAL4::VP16}A11* promote expression after zygotene, *P{GAL4-nos.NGT}A* promotes expression only in the germarium (during prophase I), and *P{GAL4: :VP16-nos.UTR}CG6325^MVD1^* promotes expression throughout the whole germline. In addition, heat shock was used to induce expression of transgenes using *P{GAL4-Hsp70,PB}89-2-1*.

The shRNA used to reduce *Cenp-C* expression, *GL00409*, is located on chromosome II. To measure the extent of the *Cenp-C mRNA* knockdown, total RNA was extracted from stage 14 oocytes expressing *GL00409* with *P{matalpha4-GAL-VP16}V37* using the TRIzol Reagent (Life Technologies). cDNA was made using the High-Capacity cDNA Reverse Transcription kit (Applied Biosystems). TaqMan Gene Expression Assays (Life Technologies) were used to measure expression levels of CENP-C. RT-qPCR was done on a StepOnePlus (Life Technologies) real-time PCR system and results showed that the *Cenp-C* RNAi expressed only 4% of wild-type RNA. In addition, CAL1 was knocked down using shRNA *GL01832* and CDK1 was knocked down using shRNA *HMS01531*, both located on chromosome II. Both shRNAs were sterile in combination with *P{matalpha4-GAL-VP16}V37* and *HMS01531* oocytes failed to enter meiosis I.

*Cencp-C^Z3-4375^* is a homozygous-viable allele that has a missense mutation in the CENP-C motif, changing a proline amino acid to a serine at position 1116 [47]. For the analysis of *Cenp-C^Z3-4375^*, trans heterozygous mutant strains were used, with one chromosome carrying either the *Cenp-C^IR35^* or *Cenp-C^pr141^* mutation. The *Cenp-C^IR35^* mutation is a premature stop codon at position 858 before the CENP-C motif and Cupin domain, resulting in a deletion of these regions [64]. The *Cenp-C^pr141^* mutation is a premature stop codon at position 1107 within the CENP-C motif [64], resulting in a partial deletion of the CENP-C motif and the rest of the protein.

### HA- and GFP-tagged transgenes

The coding region of cDNA clone FI18815 was PCR amplified and cloned in frame into pENTR4 (HiFi assembly, NEB) and then into pPHW using Clonase (Life Tech.). The target sequence for *GL00409* is located in the 5’UTR of *Cenp-C* and, therefore, this transgene is RNAi resistant. The GFP-CENP-C transgenes were constructed by Christian Lehner and contain the 5’UTR sequences upstream of the GFP sequence, and thus is RNAi sensitive. MIS12 localization was observed using a UASP regulated GFP-fusion transgene [65].

### Fertility trials, nondisjunction, and crossover assays

The *GL00409 Cenp-C* shRNA was tested for fertility and chromosome segregation errors by crossing females expressing the shRNA to *yw/ B^S^ Y* males. The aneuploid genotypes that survive are *y/y/ B^S^ Y* (Bar-eyed females) and *y w / 0* (wild-type males). Because half the aneuploid progeny have lethal genotypes, aneuploidy was calculated by multiplying the number of aneuploid progeny by two and dividing by the total number of progeny.

Sister chromatid aneuploidy was tested by crossing *y w/ Bwinscy* females expressing the shRNA to *v f B ^ Y* males. If aneuploidy occurs among homologous chromosomes, the expected genotypes that survive are *y w/Bwinscy* and *O/ v f B ^ Y*. Aneuploidy among sister chromatids is detected by the genotypes *y w/y w* (non-bar-eyed females) or *Bwinscy/Bwinscy* (Bar-eyed females). Because the O/ *v f B ^ Y* genotype arises from both MI or MII nondisjunction, only the *y w/y w* and *Bwinscy/Bwinscy* progeny were used to determine the sister chromatid aneuploidy frequency. Therefore, the number of aneuploid progeny counted was multiplied by four and divided by the total number of progeny in order to measure sister chromatid aneuploidy.

In order to observe the role of CENP-C in crossing over, females were generated that expressed the shRNA and were heterozygous for four genetic markers on chromosome III: *st, cu, e* and *ca*. These females were crossed to a strain carrying all the recessive traits and the frequency of the recombinants was scored.

### Cytology and immunofluorescence of early prophase oocytes

For immunolocalization experiments in the germaria, mated females were aged for 1-2 days at 25°C. In one well of a two well plate, 10-15 ovaries were dissected using 1x Robb’s media and then were transferred to the second well containing fresh media. A tungsten needle was used to break open the ovary sheath and tease the ovarioles apart. The dissected ovaries were transferred to a 1.5 mL Eppendorf tube with 4% formaldehyde in 500 ul of Buffer A as described [66]. The ovaries were nutated at room temperature for 10 minutes, then were washed four times before adding the primary antibodies. The next day, following four washes, secondary antibodies were added and incubated at room temperature for 4 hours.

### Cytology and immunofluorescence of metaphase I oocytes

For immunolocalization experiments in pro-metaphase I oocytes (stages 13-14), we used the immunocytochemical protocol as described [67]. In brief, 100-200 females were aged 2-3 days with males in yeasted vials [68]. Oocytes were collected by pulsing the females in a blender and then separating the oocytes from the bulk fly tissues using a mesh. Oocytes were fixed in 5% formaldehyde solution for 2.5 minutes, and then equal amounts of heptane were added and the oocytes were vortexed for 30 seconds. The membranes were removed by rolling the oocytes between a coverslip and the frosted part of a glass slide. These oocytes were incubated in PBS/1% Triton X-100 for two hours, then washed in PBS/0.05% Triton X-100. The oocytes were blocked in PBS/0.1% Tween 20/0.5% BSA (PTB) for one hour, and then incubated with primary antibodies overnight. Oocytes were washed the next day in PTB and incubated with secondary antibodies for 4 hours at room temperature.

Tissues were mounted for confocal imaging using SlowFade Gold (Invitrogen). A Leica TPS SP8 confocal microscope with a 63X, N.A. 1.4 lens was used to visualize fluorescent tags using different colored lasers. Images were imaged by collecting sections throughout the germarium or stage 14 oocyte spindle, using parameters optimized by the Leica Confocal software based on the lens and wavelength. Images were analyzed as image stacks and presented as maximum projections of whole germarium, cells, or spindles.

### Antibodies

An antibody against CENP-C made in guinea pig was made by generating a clone expressing amino acids 502-939 (Genscript). This guinea pig anti-CENP-C was used at 1:1000. Additional primary antibodies were rat anti-CID (Active motif, 1:100), rabbit anti-CID (Active motif, 1:100), rabbit anti-SPC105R (1:4000) [69], rabbit anti-GFP (Invitrogen, 1:400), rat anti-HA (Roche, 1:50), mouse anti-C(3)G (1:500) [70], two mouse anti-ORB antibodies, 6H4 and 4H8 (1:100 for each) [71] and mouse anti-α tubulin DM1A conjugated directly to FITC (Sigma, 1:50). The secondary antibodies that were used were Cy3, Alexa 546, Alexa 633, or Alexa 647 from Jackson Immunoresearch Laboratories, and Alexa 488 from Invitrogen. The oocytes were stained with Hoechst 33342 at 1:10,000 (10 μg/ml). FISH probes were obtained from IDT where the X359 repeat was labeled with Alexa 594, the dodeca repeat was labeled with Cy5, and the AACAC repeat was labeled with Cy3.

### Quantification and statistical analysis

Aneuploidy in flies expressing the *Cenp-C* shRNA with NGTA was compared to that of the control group and a t-test was done to determine if the difference is statistically significant (p-value < 0.05). Sister chromatid aneuploidy in flies expressing the *Cenp-C* shRNA with NGTA was compared to that of the control group and a t-test was used to determine whether the difference was statistically significant (p-value < 0.05). The percent of crossovers at each position was compared to that of the control group using a t-test, and if the difference was statistically significant (p-value < 0.05), then this indicated that elevated or reduced recombination occurred at that specific location.

Quantification of CID foci and protein localization in both the germarium and in stage 14 oocytes was measured using the Imaris Software. To quantify centromere foci, the automated spots detection feature of Imaris was used. A spot with a XY diameter of 0.20 μm, a Z diameter of 1.00 μm, and a physical interaction with the DNA was counted as a centromere. A t-test was done to compare the number of centromere foci in flies expressing the knockdown to the number in the control group using a p-value < 0.05 to indicate statistical significance.

To measure the localization of proteins to the centromere or the nucleus in *Cenp-C* RNAi, intensity experiments were performed using Imaris. The Imaris software was utilized to measure protein intensity at the centromere of the oocyte, using CID as the centromere marker. Spots were chosen in random somatic cells to quantify the intensity of the background. The intensity of the protein at the centromere was divided by the background and a t-test was used to determine whether the centromere protein intensity was reduced with a loss of CENP-C.

Intensity experiments were also performed to measure localization of MIS12, SPC105R, or CENP-C to the centromere at stage 14 using a similar protocol. However, background intensity was measured in the oocyte cytoplasm as opposed to measuring the background in the somatic cells for germarium images. A t-test was used to determine whether the intensity of the protein of interest at the centromere normalized to the background was reduced with a loss of CENP-C using a p-value <0.05 to indicate statistical significance.

## Supplementary Figures

Figure S 1: **CAL1 and CID localization in oocytes**. A) Localization of GFP-CID (green) and centromeres detected using a CENP-C antibody (red). B) Localization of GFP-CAL1 (green) with the centromeres detected using a CID antibody (red). (D) Localization of GFP-CENP-C, with the centromeres detected using a CID antibody (red). In all images, the DNA is blue and the scale bars are 5 mm.

Figure S 2: **GFP-CENP-C, CID-GFP or CAL1-GFP in stage 14 oocytes.** The DNA is blue, microtubules in white, and the scale bars represent 5 um. **A)** Localization of CID-GFP or CAL1-GFP (green) using the *MVD1, mata* or *NGTA*. In the one image (*gcal1-GFP)*, CAL1-GFP is regulated by the endogenous *cal1* promoter. The centromeres were detected using CENP-C (red). **B)** Localization of CID-GFP or GFP-CENP-C (green) using *mata*. The kinetochores were detected using an antibody against SPC105R (red).

Figure S 3: Expression pattern of NGTA. The expression pattern of **(A)** *P{GAL4-nos.NGT}* and **(B)** *P{GAL4::VP16-nos.UTR}CG6325MVD1* using *UASP-β-galactosidase* as a reporter. Arrowheads indicate anterior tip of the ovariole, where the germarium is located, and the blue stain indicates where each GAL4 promotes expression.

Figure S 4: Loading of HA-CENP-C during oocyte meiotic prophase. In all images, HA-CENP-C is green, the centromeres are marked with CID (red), and DNA is in blue. The scale bars represent 5 um. **A)** Whole germarium with HA-tagged CENP-C expressed using *NGTA* or *mata*. ORB (white) is enriched in the oocyte. **B)** HA-tagged CENP-C or CENP-C^C^ was expressed using *mata*. **C)** HA-CENP-C was expressed using *hsp70-Gal4*. Oocytes were collected and fixed after 6 hours (early prophase) or 5 hours (stage 14) after a 1-hour incubation at 37°C.

Figure S 5: SC assembly when CENP-C is depleted in prophase. Confocal images of the germarium with *Cenp-C* RNAi (HMS01171) with (A) no *GAL4* and (B) *NGTA*. DNA is shown in blue, CENP-C is in red, and C(3)G is in green. The scale bar is 10 µm. CENP-C and C(3)G are shown in white in the single channel images. Region 1 of the germarium has been boxed to show increased centromeric C(3)G. The insets show single nuclei from region 1 in the germarium to show co-localization of CENP-C and C(3)G (Scale bar= 3 µm).

Figure S 6: Additional images of CENP-C localization in RNAi and transgenic oocytes. **A)** *Cenp-C* RNAi or *Cenp-C^Z^*oocytes with CENP-C (green) and CID (red). **B)** *Cenp-C* RNAi or *Cenp-C^Z^* oocytes expressing a *Cenp-C* transgene, with HA in red and SPC105R in green, DNA in blue, microtubules in white, and the scale bars represent 5 um. **C)** Oocytes shown in panels A and B were assessed for KT-MT attachments. This was done by measuring the distance between each centromere and the nearest microtubule. **D)** Number of centromere foci was measured based on CID foci in *Cenp-C* RNAi metaphase I oocytes (n = 19 and 36). Error bars represent standard deviation.

Figure S 7: Loading during prophase does not depend on CDK1 or CAL1. Localization of HA-CENP-C in stage 5 oocytes of **(A)** *cdk1* and **(B)** *cal1* RNAi oocytes. HA is in red, CID is in white, cytoplasmic ORB protein is in green, and DNA is in blue. The scale bar represents 5 um. Relative intensity of CENP-C in control and RNAi oocytes (n=10, 12, 11).

