## Supplemental Figures for "A Dynamic population of prophase CENP-C is required for meiotic chromosome segregation"

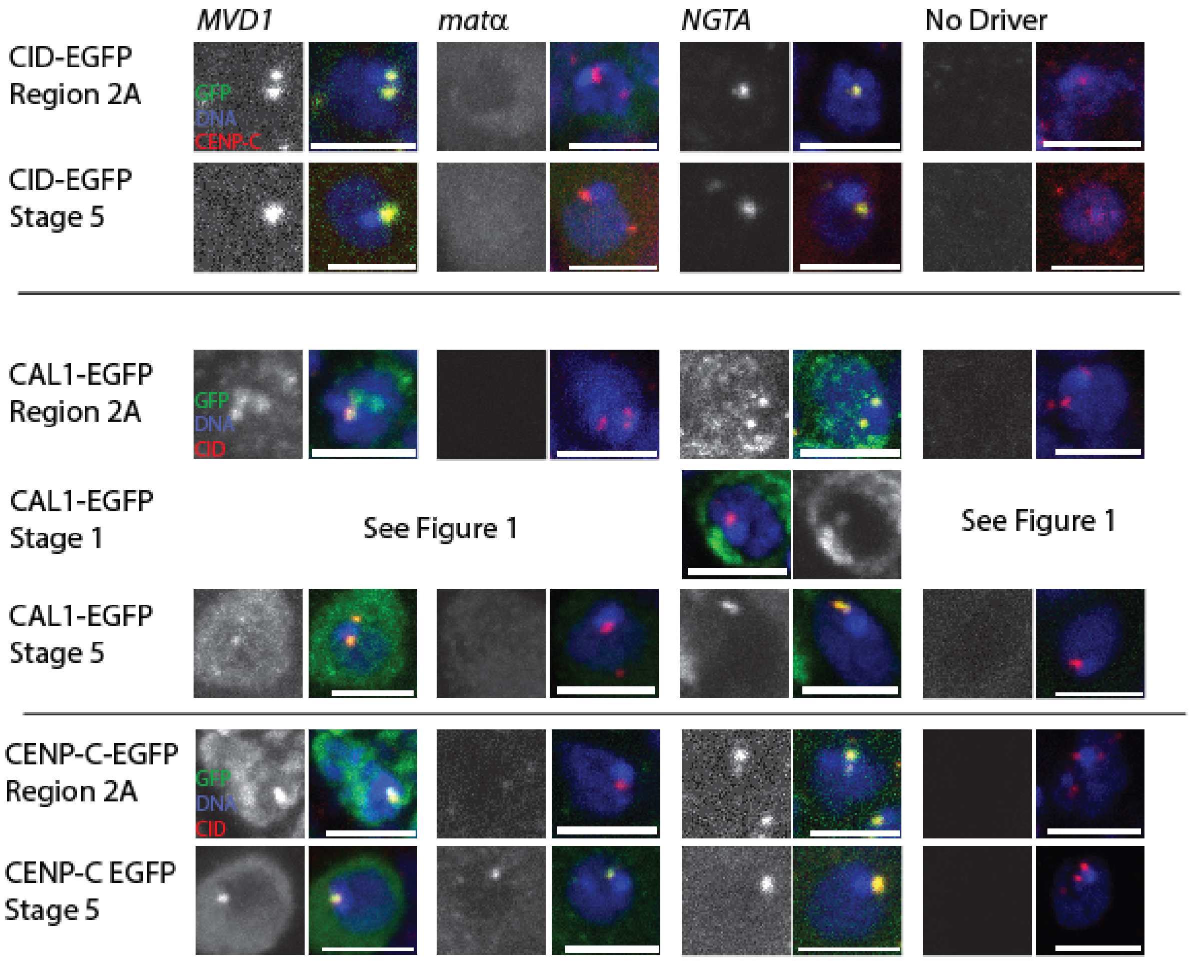


Figure S 1: **CAL1 and CID localization in oocytes**. A) Localization of GFP-CID (green) and centromeres detected using a CENP-C antibody (red). B) Localization of GFP-CAL1 (green) with the centromeres detected using a CID antibody (red). (D) Localization of GFP-CENP-C, with the centromeres detected using a CID antibody (red). In all images, the DNA is blue and the scale bars are 5 mm.

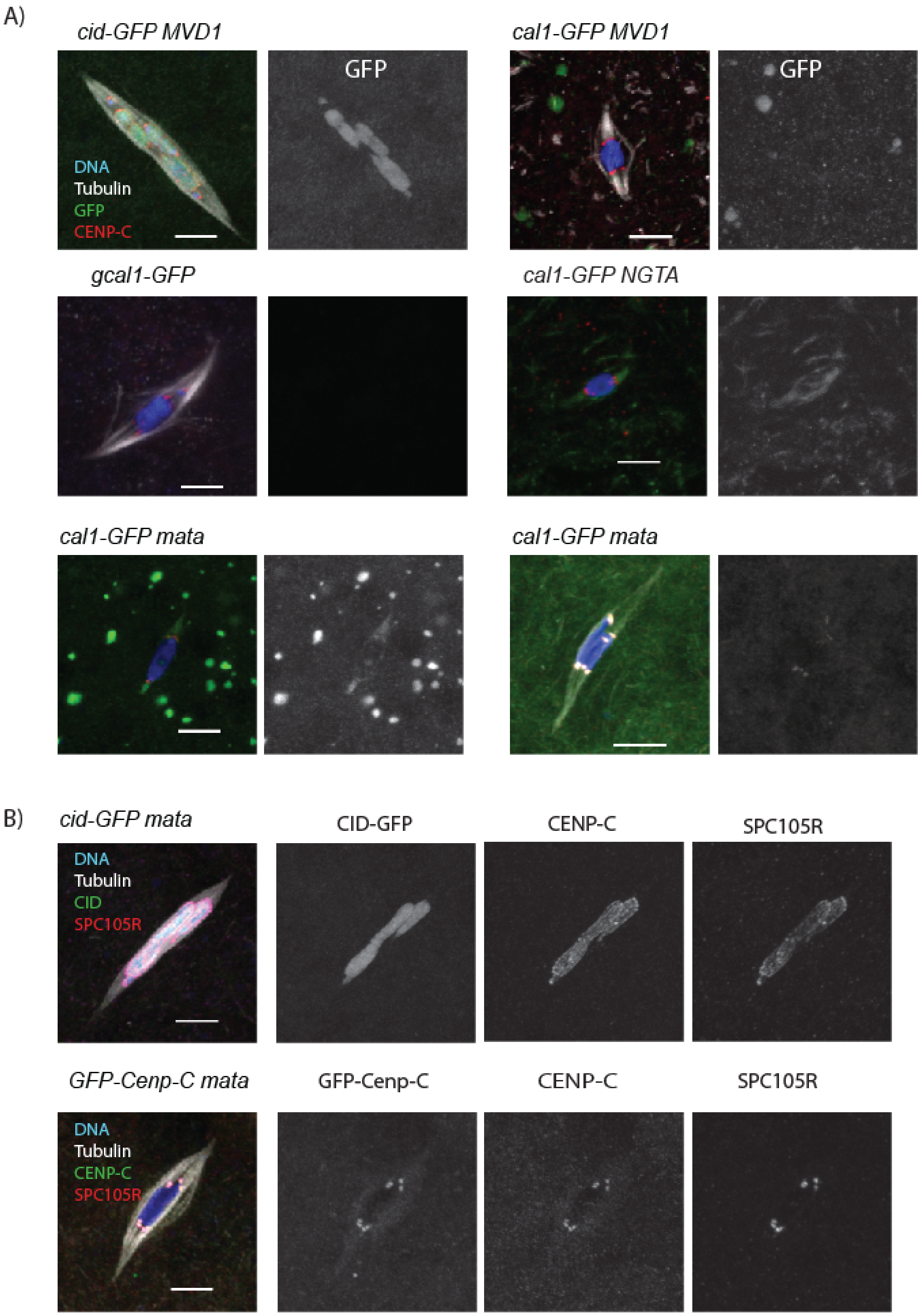


Figure S 2: **GFP-CENP-C, CID-GFP or CAL1-GFP in stage 14 oocytes.** The DNA is blue, microtubules in white, and the scale bars represent 5 um. **A)** Localization of CID-GFP or CAL1-GFP (green) using th e *MVD1, mata* or *NGTA*. In one image, CAL1-GFP is regulated by the endogenous *cal1* promoter. The centromeres were detected using CENP-C (red). **B)** Localization of CID-GFP or GFP-CENP-C (green) using *mata*. The kinetochores were detected using an antibody against SPC105R (red).

A) *P{GAL4-nos.NGT}A*

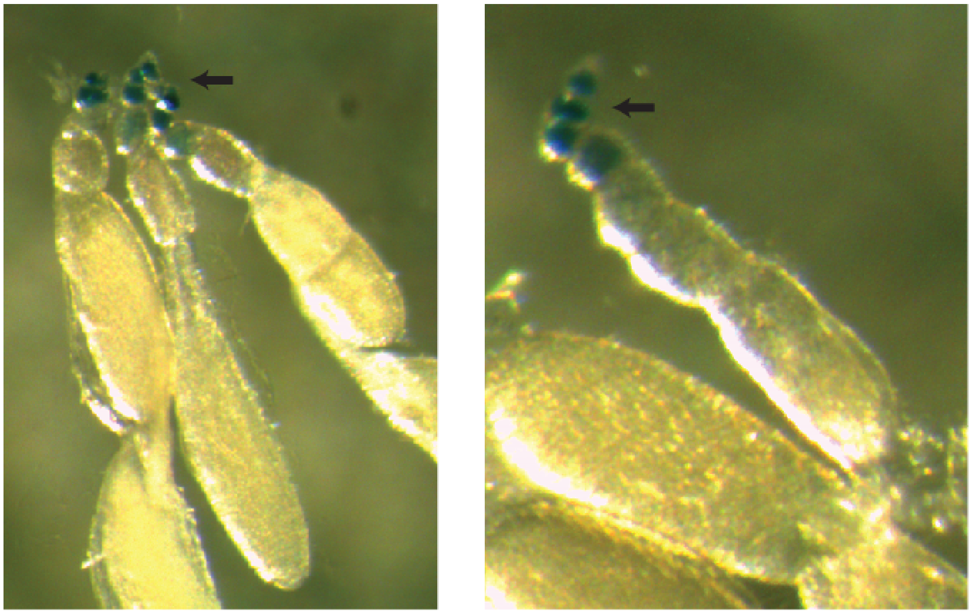

B) *P{GAL4: :VP16-nos.UTR}CG6325MVD1*

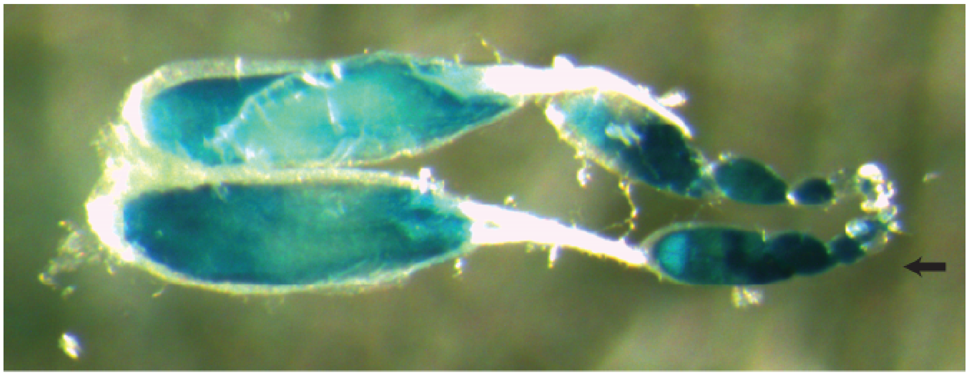

Figure S 3: Expression pattern of NGTA. The expression pattern of **(A)** *P{GAL4-nos.NGT}* and **(B)** *P{GAL4::VP16-nos.UTR}CG6325MVD1* using *UASP-β-galactosidase* as a reporter. Arrowheads indicate anterior tip of the ovariole, where the germarium is located, and the blue stain indicates where each GAL4 promotes expression.

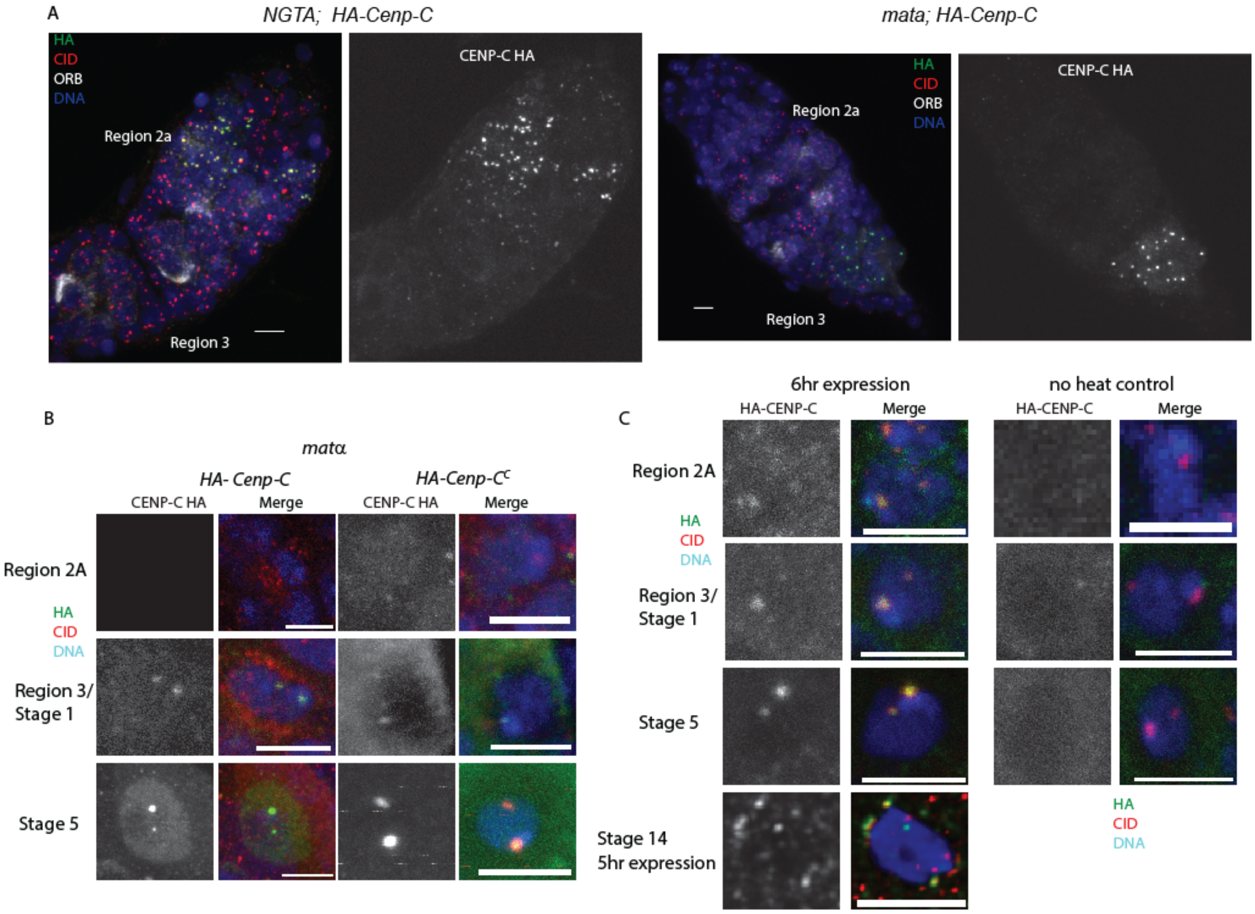


Figure S 4: Loading of HA-CENP-C during oocyte meiotic prophase. In all images, HA-CENP-C is green, the centromeres are marked with CID (red), and DNA is in blue. The scale bars represent 5 um. **A)** Whole germarium with HA-tagged CENP-C expressed using *NGTA* or *mata*. ORB (white) is enriched in the oocyte. **B)** HA-tagged CENP-C or CENP-C$^{C}$ was expressed using *mata*. **C)** HA-CENP-C was expressed using *hsp70-Gal4*. Oocytes were collected and fixed after 6 hours (early prophase) or 5 hours (stage 14) after a 1-hour incubation at 37°C.

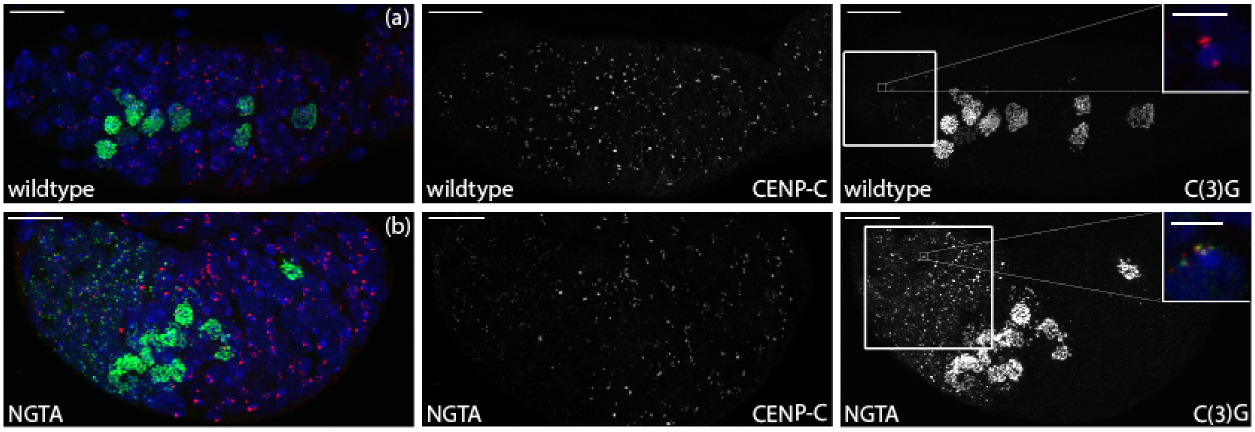


Figure S 5: SC assembly when CENP-C is depleted in prophase.  Confocal images of the germarium with *Cenp-C* RNAi (HMS01171) with (A) no *GAL4* and (B) *NGTA*. DNA is shown in blue, CENP-C is in red, and C(3)G is in green. The scale bar is 10 μm. CENP-C and C(3)G are shown in white in the single channel images. Region 1 of the germarium has been boxed to show increased centromeric C(3)G. The insets show single nuclei from region 1 in the germarium to show co-localization of CENP-C and C(3)G (Scale bar= 3 μm).

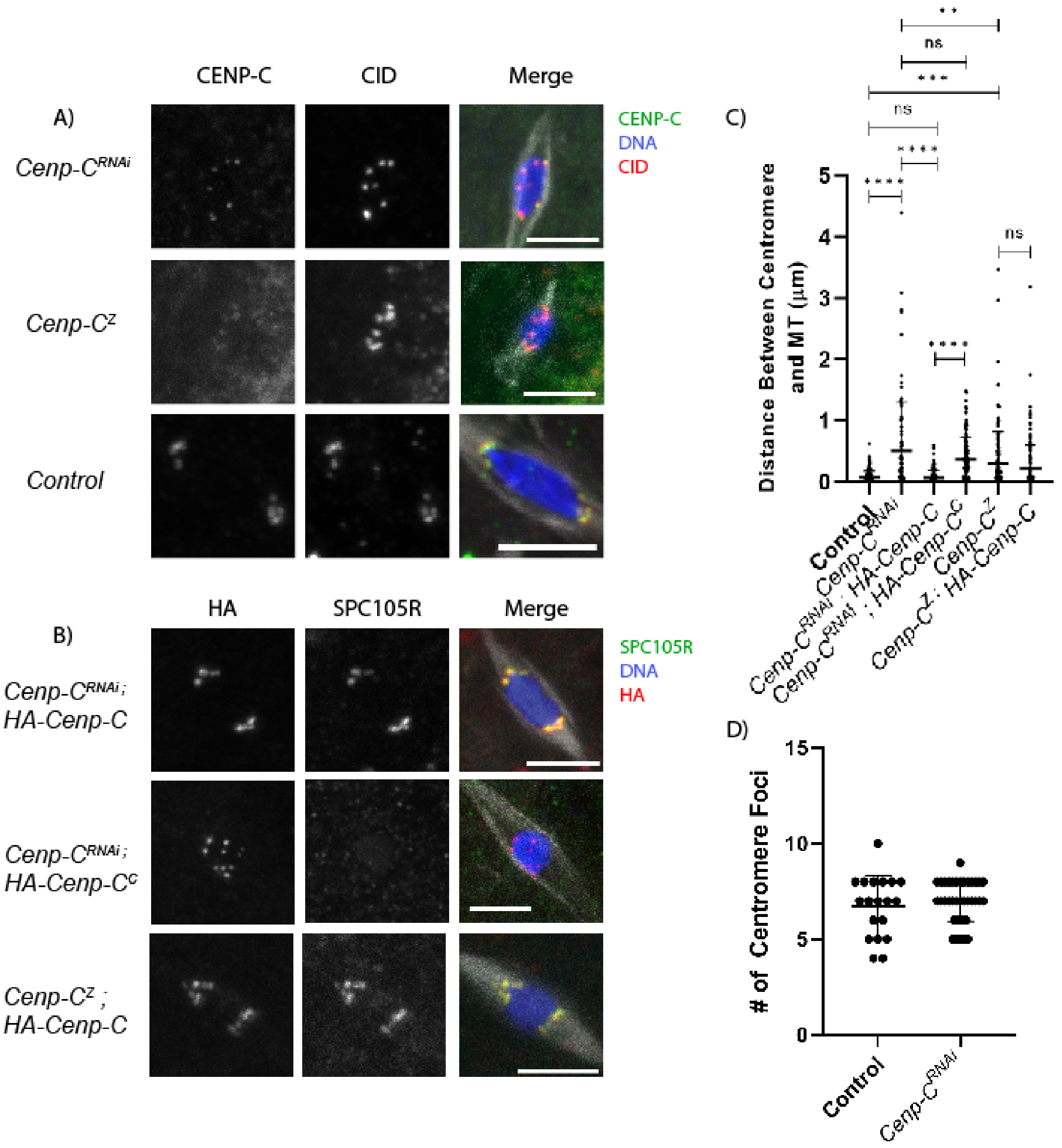


Figure S 6: Additional images of CENP-C localization in RNAi and transgenic oocytes. **A)** *Cenp-C* RNAi or *Cenp-C*$^{Z}$ oocytes with CENP-C (green) and CID (red). **B)** *Cenp-C* RNAi or *Cenp-C*$^{Z}$ oocytes expressing a *Cenp-C* transgene, with HA in red and SPC105R in green, DNA in blue, microtubules in white, and the scale bars represent 5 um. **C)** Oocytes shown in panels A and B were assessed for KT-MT attachments. This was done by measuring the distance between each centromere and the nearest microtubule. **D)** Number of centromere foci was measured based on CID foci in *Cenp-C* RNAi metaphase I oocytes (n = 19 and 36). Error bars represent standard deviation.

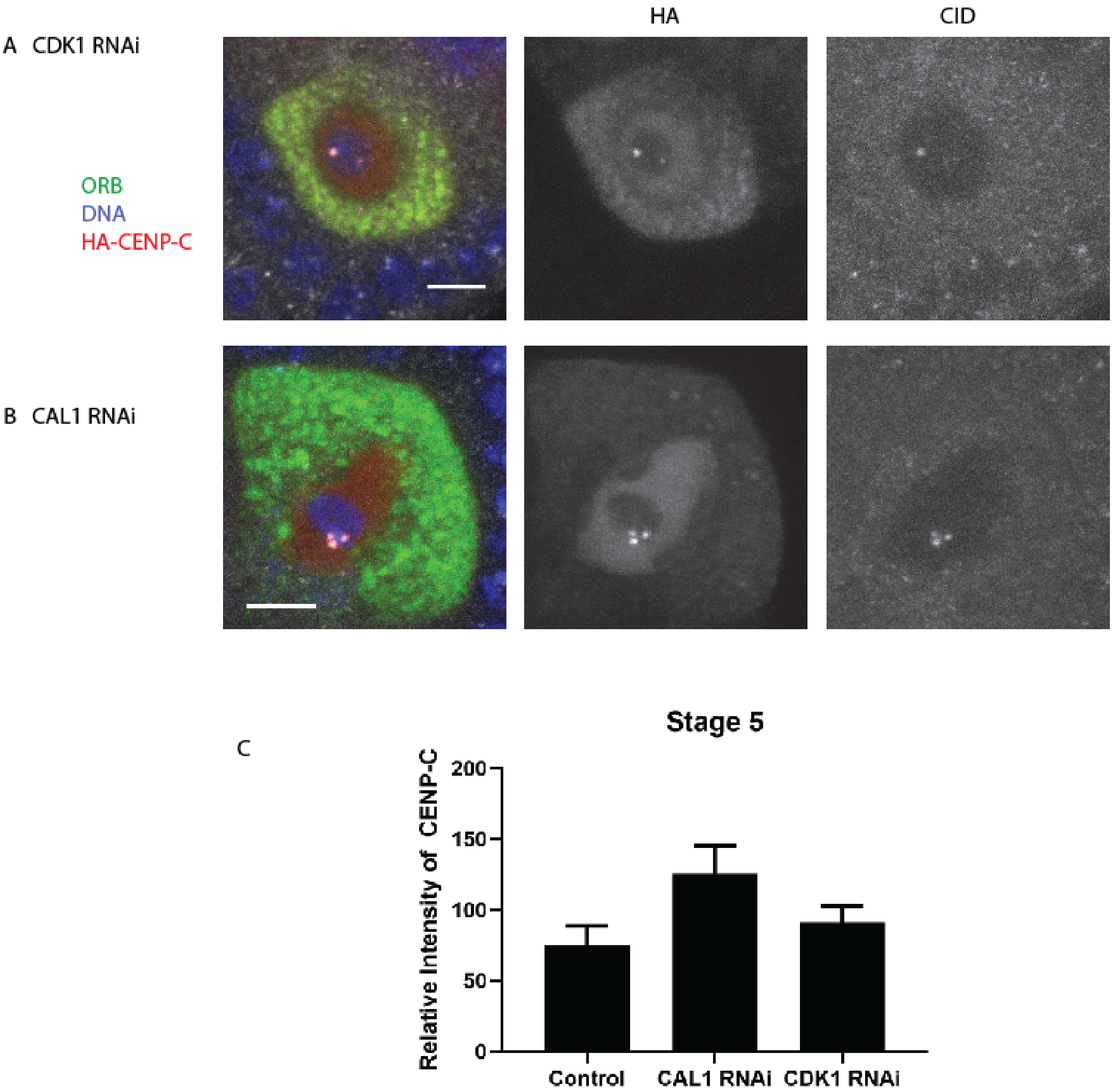


Figure S 7: Loading during prophase does not depend on CDK1 or CAL1. Localization of HA-CENP-C in stage 5 oocytes of **(A)** *cdk1* and **(B)** *cal1* RNAi oocytes. HA is in red, CID is in white, cytoplasmic ORB protein is in green, and DNA is in blue. The scale bar represents 5 um. **C)** Relative intensity of CENP-C in control and RNAi oocytes (n=10,12, 11).
